# E-cadherin interacts with EGFR resulting in hyper-activation of ERK in multiple models of breast cancer

**DOI:** 10.1101/2020.11.04.368746

**Authors:** Gabriella C. Russo, Ashleigh J. Crawford, David Clark, Julie Cui, Ryan Carney, Michelle N. Karl, Boyang Su, Bartholomew Starich, Tung-Shing Lih, Pratik Kamat, Qiming Zhang, Pei-Hsun Wu, Meng-Horng Lee, Hon S. Leong, Vito W. Rebecca, Hui Zhang, Denis Wirtz

**Affiliations:** Department of Chemical and Biomolecular Engineering, Johns Hopkins University, 3400 N Charles St, Baltimore, Maryland 21218, USA; Johns Hopkins Physical Sciences– Oncology Center and Institute for NanoBioTechnology, Johns Hopkins University, 3400 N Charles St, Baltimore, Maryland 21218, USA; Department of Pathology, The Johns Hopkins University School of Medicine, Baltimore, Maryland 21231, USA; Department of Biophysics, Johns Hopkins University, 3400 N Charles St, Baltimore. MD 21218, USA; Department of Medical Biophysics, University of Toronto, Toronto, ON; Biological Sciences Platform, Sunnybrook Research Institute, Toronto, ON; Department of Biochemistry and Molecular Biology, Johns Hopkins University School of Public Health, Baltimore, MD 21231, USA

## Abstract

The loss of the intercellular adhesion molecule E-cadherin is a hallmark of the epithelial-mesenchymal transition (EMT), during which tumor cells transition into an invasive phenotype. Accordingly, E-cadherin has long been considered a tumor suppressor gene. Using novel multi-compartment spheroids and multiple *in vivo* models, we show that E-cadherin promotes a hyper-proliferative phenotype in breast cancer cells via interaction with the transmembrane receptor EGFR. This interaction results in the activation of the MEK/ERK signaling pathway, leading to a significant increase in proliferation via the activation of transcription factors including c-Fos. Pharmacological inhibition of MEK activity in E-cadherin positive breast cancer cells significantly decreases both tumor growth and macro-metastasis *in vivo*. This work provides evidence for a novel role of E-cadherin in breast tumor growth and identifies a potential new target to treat hyper-proliferative E-cadherin-positive breast tumors.

## INTRODUCTION

E-cadherin (E-cad), a transmembrane molecule ubiquitously expressed in normal epithelial tissues, promotes and maintains intercellular adhesion. In cancer cells, the loss of E-cad expression and gain of N-cadherin expression is associated with onset of invasion via the epithelial-to-mesenchymal transition (EMT) process.^1, 2^ EMT consists of highly orchestrated cascade of molecular events where epithelial cells transition from a polarized, stationary state to an invasive, migratory phenotype that is accompanied by changes in cell morphology.^1, 2^ These processes are believed to then trigger metastasis in carcinomas (cancers of epithelial origin). Together these results have long suggested that E-cad is a tumor suppressor gene and that its expression correlates with a better survival outcome in patients.^1, 2^ However, considering that the most common type of breast cancer, invasive ductal carcinoma (IDC), is mostly E-cad positive and can retain E-cad expression when metastasizing, its classification as a tumor suppressor gene is being re-assessed.^3, 4^

Clinical data indicate that high E-cad expression in IDC is associated with a worse overall survival rate of patients.^4, 5^ This suggests that E-cad may function as an oncogene in breast cancer rather than a tumor suppressor gene. Recent studies aimed at re-assessing E-cad function in cancer have focused on the role of E-cad in EMT^11–14^. Whether and how E-cad expression impacts proliferation in cancer cells is not understood and the functional implications of E-cad interactions with other transmembrane proteins, like EGFR, have yet to be fully elucidated^6–10^. E-cad is activated by dual functioning β-catenin at the cell membrane^15, 16^. To activate E-cad, β-catenin forms a complex with α-catenin and binds to the cytoplasmic tail of E-cad; once activated, E-cad links to the actin cytoskeleton and can form adherens junctions between neighboring cells. When not interacting with E-cad, β-catenin can also function as a transcription factor in the nucleus where it can impact cell growth^15, 16^. E-cad has also been shown to interact with and subsequently phosphorylate EGFR in normal epithelial cells, which can activate proliferative pathways^10, 17^. Whether E-cad/EGFR interactions occur in cancer cells and how their interactions impact tumor progression, including cancer-cell proliferation, remain unexplored^6–8^. Thus, it is important to examine the role of E-cad in breast cancer cell proliferation to further understand the apparent disparity between the clinical data and our understanding of how E-cad functions in cancer.

Here, utilizing new multi-compartment spheroids and multiple *in vivo* models, we studied the impact of E-cad expression on breast cancer cell proliferation, at both the primary and secondary metastatic sites *in vivo.* Remarkably, E-cad expression in breast cancer cells upregulates multiple proliferation pathways, in particular the ERK cascade within the greater MAPK signaling pathway. Through interactions with EGFR at the cell surface, E-cad expression leads to ERK hyper-activation. This E-cad-induced upregulation of the ERK cascade results in a dramatic increase in cancer cell proliferation, tumor growth and metastatic outgrowth *in vivo*. When the phosphorylation of ERK is blocked utilizing a highly specific MEK1/2 inhibitor^18^, this effect is reversed. In sum, this work suggests that E-cad plays a pro-tumorigenic role and reveals new molecular targets for patients with E-cad positive tumors, particularly invasive ductal carcinoma.

## MAIN

### E-cad expression promotes proliferation in breast cancer cells

Using Metabric^5^, we first evaluated clinical outcomes of patients with invasive ductal carcinoma (IDC), which often express high-levels of E-cadherin (E-cad). This analysis revealed that high E-cad expression is associated with a worse survival (Figure 1a)^5^. This unanticipated association between clinical outcome and E-cad expression has prompted re-evaluation of its role in EMT and metastatic disease^11–14^. Our hypothesis is that E-cad induces proliferation in E-cad+ breast cancer, further worsening the clinical outcome of patients with E-cad+ tumors. To mechanistically test this hypothesis, we first performed direct comparisons of cancer cells with high and low E-cad expression through use of engineered cell lines created from the same parental cells.

**Figure 1:**
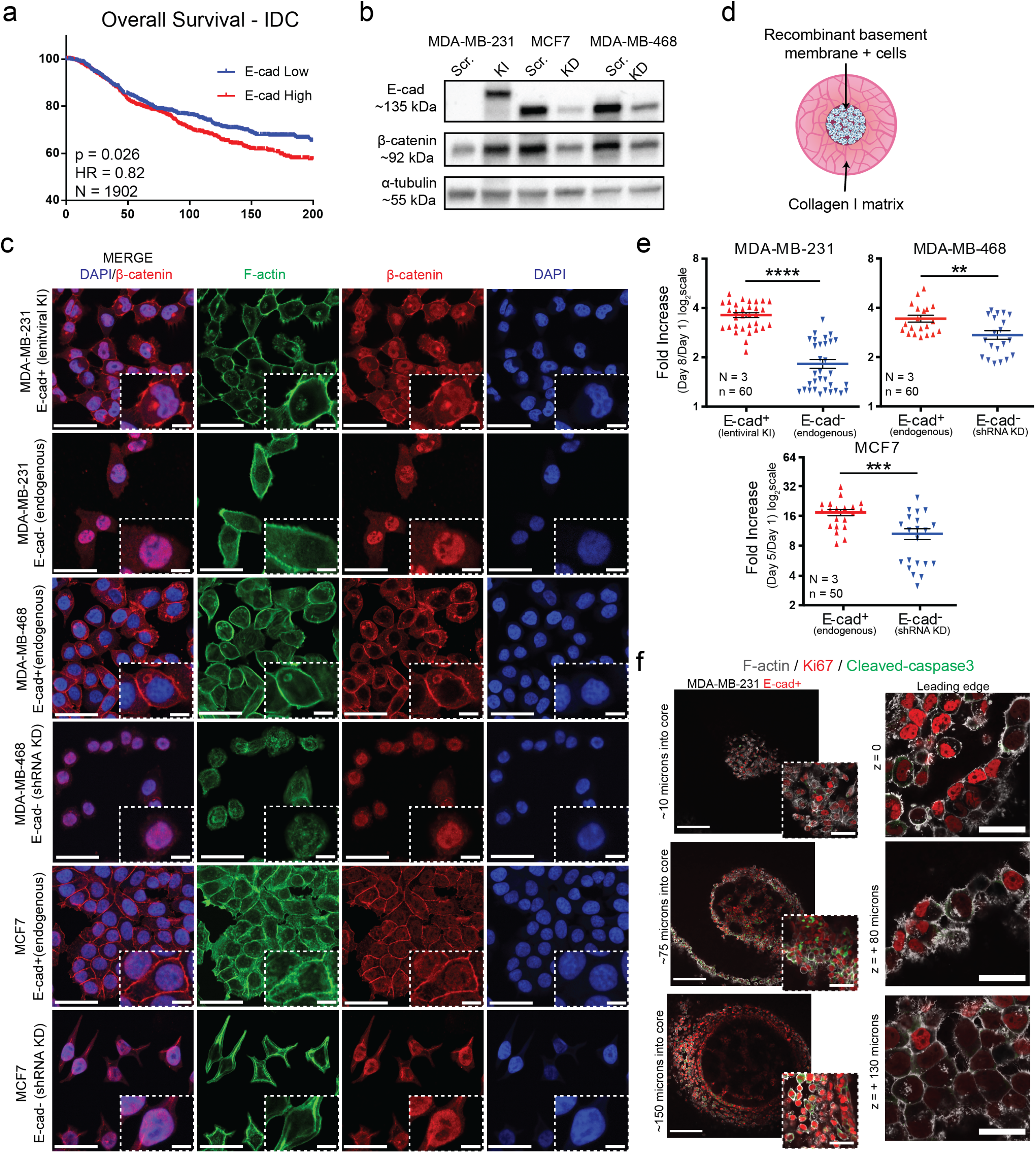
Manipulation of E-cadherin expression impacts proliferation in breast cancer cells. (a) Survival plot based on Metabric data set.^5^ Low E-cad expression is defined here as 50 percentile or lower, while high E-cad expression is 50 percentile or higher. Sample size: N =1902 patients. (b) Western blot assessment of E-cadherin, β-catenin, and a-tubulin (loading control) protein expression in E-cadherin shRNA-based knockdown (KD) or lentiviral knock-in (KI) and scramble controls (Scr.) for MDA-MB- 231, MCF7, and MDA-MB-468 human breast cancer cells. (c) Immunofluorescence micrographs showing E-cad functional status via β-catenin localization in control and E-cad modified MDA-MB-231, MDA-MB-468, MCF7 cells *in vitro*. β-catenin is in red, F-actin actin is green, and nuclear DNA (DAPI) is blue. Scale bars = 50 µm and inset = 10 µm. N = 3. (d) Schematic of the 3D spheroid system composed of Matrigel in the inner core and collagen I outer layer. (e) PrestoBlue fold increase of MDA-MB-231 (n= 60) ****P=<0.0001, MDA-MB-468 (n= 60) **P=0.0073, and MCF7 (n= 50) ***P = 0.0008 comparing spheroids on day 1 to day 8 or day 1 to day 5. N =3 biological repeats for all cell lines. All data are mean ± SEM. Statistical test used: Mann-Whitney test, two sided for spheroids. (f) Ki67 and cleaved-caspase 3 staining of MDA-MB-231 E-cad+ spheroids on day 5 of culture. Scale bars: 200 µm, inset 50 µm and “leading edge” images scale bars: 50 µm. N =1.

We determined the endogenous expression level of E-cad in seven commonly used ductal carcinoma breast cancer cell lines, spanning all hormone receptor subtypes- HR+, HER2+, and TNBC^19^ (Extended Data Figure 1a). Of these, we chose three cell lines classified as IDC that were also suitable for *in vivo* modeling. MCF7 and MDA-MB-468 endogenously express E-cad (denoted E-cad+), while MDA-MB-231 known for its high proliferative index and metastatic potential^20, 21^ does not endogenously express E-cad (E-cad-). We generated E-cad shRNA lentiviral knockdowns for MDA-MB-468 and MCF7 cells (denoted E-cad KD) and an E-cad lentiviral knock-in for MDA-MB-231 cells (denoted E-cad KI). We performed western blotting to confirm E-cad expression changes after genetic modification (Figure 1b). We also confirmed expected functional changes associated with E-cad by assessing the -catenin within the cell. β catenin links the cytoplasmic domain of E-cad to the actin filament network, allowing E- cad to function as a cell-cell adhesion molecule^17^. Immunofluorescence microscopy showed that β-catenin accumulated in the nucleus in E-cad- cells, as anticipated, and localized in the cytoplasm and at the cell membrane of E-cad+ cells, where β atenin can function in conjunction with E-cad^22^ (Figure 1c). As anticipated^22, 23^, we also observed organized actin filaments in E-cad+ cells and a more disorganized, wavy appearance in E-cad- cells (Figure 1c).

We then compared the proliferation of these cell pairs in standard culture conditions and observed a significantly higher proliferation rate of E-cad+ cells compared to E-cad- cells (Extended Data Figure 1b). However, this 2D environment does not take into account the 3D nature of a tumor and cells are artificially apically polarized. Hence, we developed a 3D double-layer spheroid system that better mimics the primary tumor environment to study the impact of E-cad on cancer cell proliferation. The core of these spheroids contains a controlled number of cancer cells suspended in Matrigel, which is then encapsulated in a collagen I matrix, resulting in a collagen corona around the Matrigel core of cancer cells. This model mimics the basement membrane and the surrounding collagen-rich stroma of solid breast tumors (Figure 1d) and allows us to monitor the proliferation and invasion of cancer cells that break out from the core and invade into the collagen layer.

We assessed proliferation in this spheroid model using the PrestoBlue viability assay^24^ and observed a significant increase in growth in E-cad+ spheroids compared to Ecad- spheroids (Figure 1e). We confirmed a linear relationship between RFU and cell number to ensure these results indicated increasing cell numbers (Extended Data Figure 1c). We also tracked spheroid progression via DIC imaging and observed expected^1, 2, 22, 23^ migratory changes when E-cad expression was manipulated (Extended Data Figure 1d). After 5 days of growth, we fixed and conducted immunofluorescent staining on the spheroids for proliferation marker Ki67 (red) and apoptosis marker cleaved caspase-3 (green) (Figure 1f). We observe areas of highly proliferative, Ki67+ cells in E-cad+ spheroids accompanied by some apoptosis in the core and immediate invasive layer, where cancer cells were densely packed (Figure 1f). These proliferative cells extended to the leading edge of the invasive cell front in E-cad+ spheroids as well. We did not observe any Ki67+ cells in E-cad- spheroids (Extended Data Figure 1e). These results confirm our previous observations: E-cad expression correlates with proliferation in breast cancer cells.

### E-cad expression results in hyper-activation of ERK

To understand how E-cad promoted proliferation in cancer cells, we first performed RT-qPCR analysis of genes associated with proliferation. We chose to assess two signaling pathways, one related to proliferation- the MAPK pathway- and one related to E-cad function- the WNT pathway ^25, 26^. The MAPK pathway is a three-tiered kinase cascade consisting of three main complexes – ERK, JNK, and p38. The ERK complex is of particular interest, as once it is activated, it can regulate processes such as proliferation, differentiation, and survival^25^. β-catenin is the central molecule of the WNT signaling pathway, playing roles as both a membrane-bound protein when activating E-cad and as a transcription factor when localized to the nucleus, where it can impact and regulate the cell cycle^26^. Our results show that 30% of key genes in the MAPK pathway were expressed at least 2-fold higher in E-cad+ cells compared to E-cad- breast cancer cells (Figure 2a and Extended Figure 2a). Upregulated genes in the ERK cascade included *MAP2K1 & MAP2K2* (which encode MEK1/2), *MAPK3 & MAPK1* (which encode ERK1/2), and *FOS* (downstream transcription factor). In the WNT signaling pathway, approximately 27% of genes were expressed at least 2-fold higher when E-cad was present (Figure 2a). In addition to the upregulation of β-catenin (*CTNNB1*), we observed upregulation of *GSK3*β (Figure 2a), which can degrade β-catenin through interactions with *APC* and *AXIN1*^26^. Genes downstream of *CTNNB1*, including Lymphoid Enhancer Binding Factor 1 (*LEF1*) and Cyclin D1 (*CCND1*) were concurrently upregulated.

**Figure 2:**
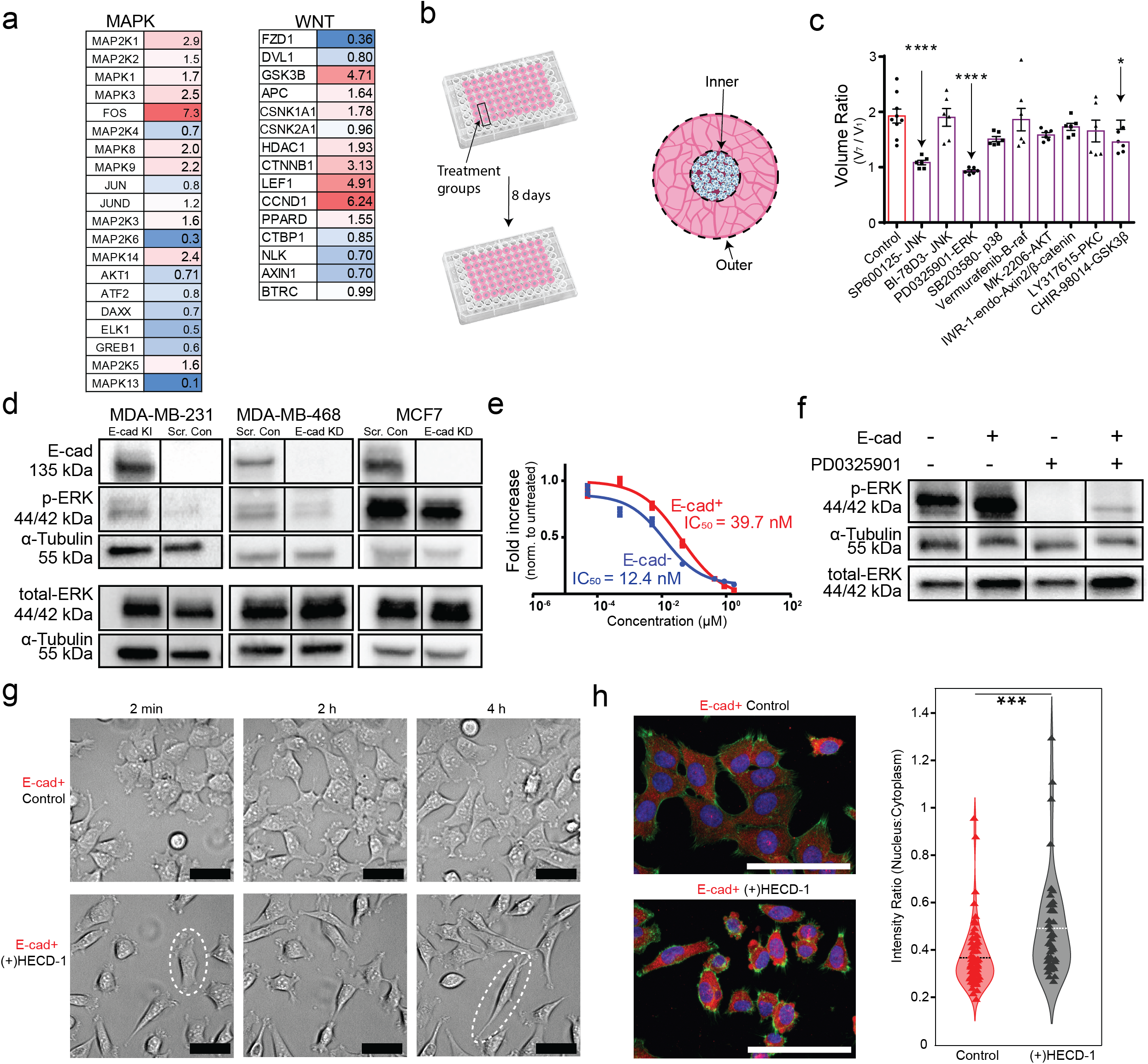
E-cadherin promotes hyper-proliferation via the MAPK pathway. (a) Relative expression of genes in the MAPK and WNT signaling pathways in MDA-MB- 231 E-cad+ to MDA-MB-231 E-cad- cells assessed via RT-qPCR. Fold change of +/- 2 is considered biologically relevant. N=3 biological repeats. (b) Schematic of high- throughput experiment to assess response to inhibitors listed in Table 1 and schematic of volume measurements used to calculate inner and outer volume ratios. (c) Inner volume ratio of 10 µM inhibitor screening (N=1, n=5-6 per condition). (d) IC_50_ of PD0325901 E-cad+ cells require 4x more inhibitor to achieve the same effect in E-cad- cells. p value <0.0001. (e) Representative western blot results, confirming higher levels of p-ERK in E-cad+ spheroids for all cell line pairs. (f) Western blot of MDA-MB-231 spheroids treated with MEK inhibitor PD0325901, resulting in decrease in p-ERK expression. (g) Phase contrast microscopy images of MDA-MB-231 E-cad+ during t = 2 min, 2 h, and 4 h of migration study to test impact of an E-cad functional blocking antibody. Scale bars = 50 µm. (h) Immunofluorescence staining of phosphorylated ERK (red), shown with nuclear stain (blue) and f-actin stain (green). Scale bars = 50 µm. Ratio of nuclear expression to cytoplasmic expression of phospho-ERK increases in response to HECD-1 treatment, indicating more phospho-ERK is localizing to the nucleus. N = 3 biological repeats, n = 4 technical repeats.

**Table 1:**
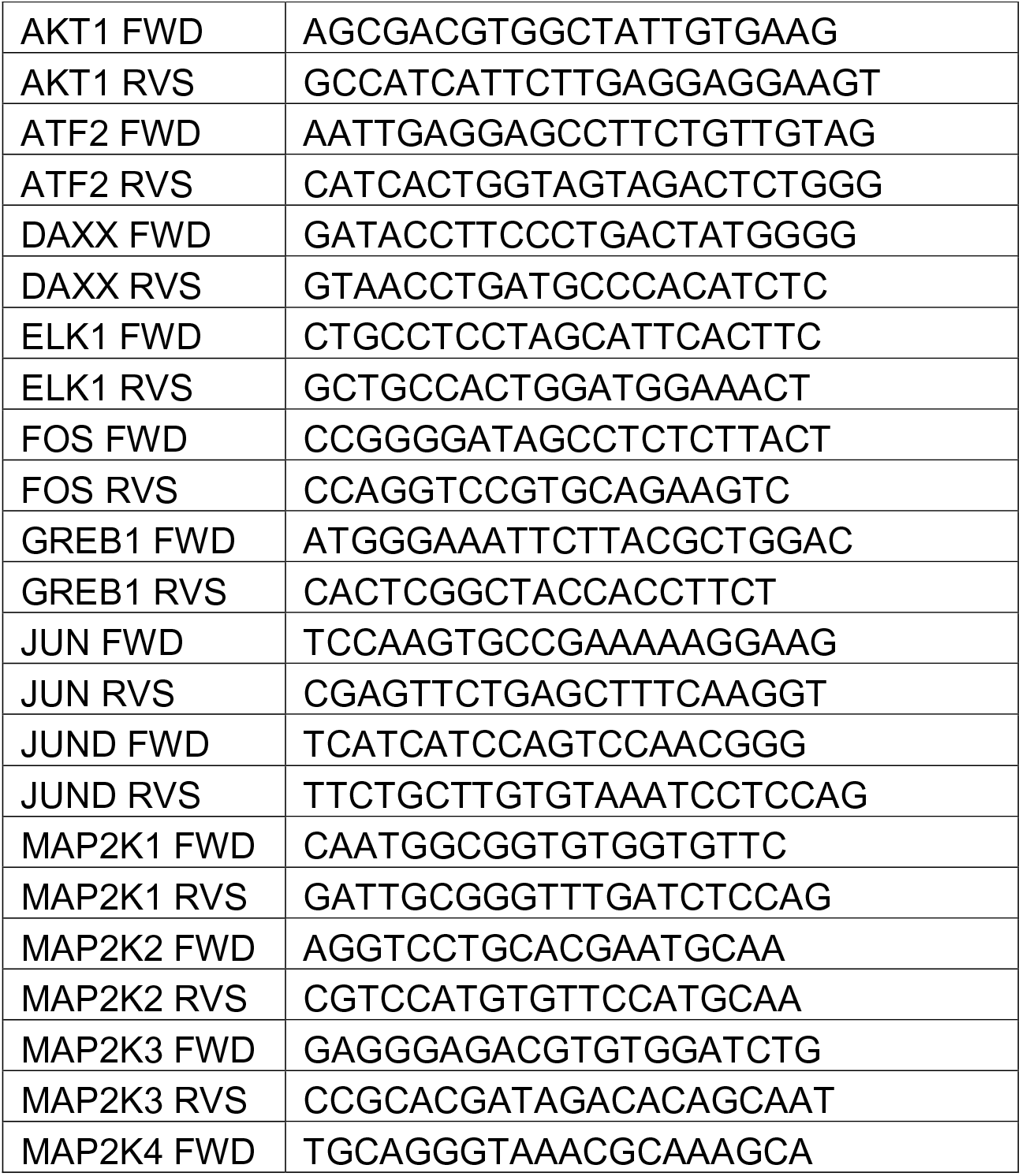
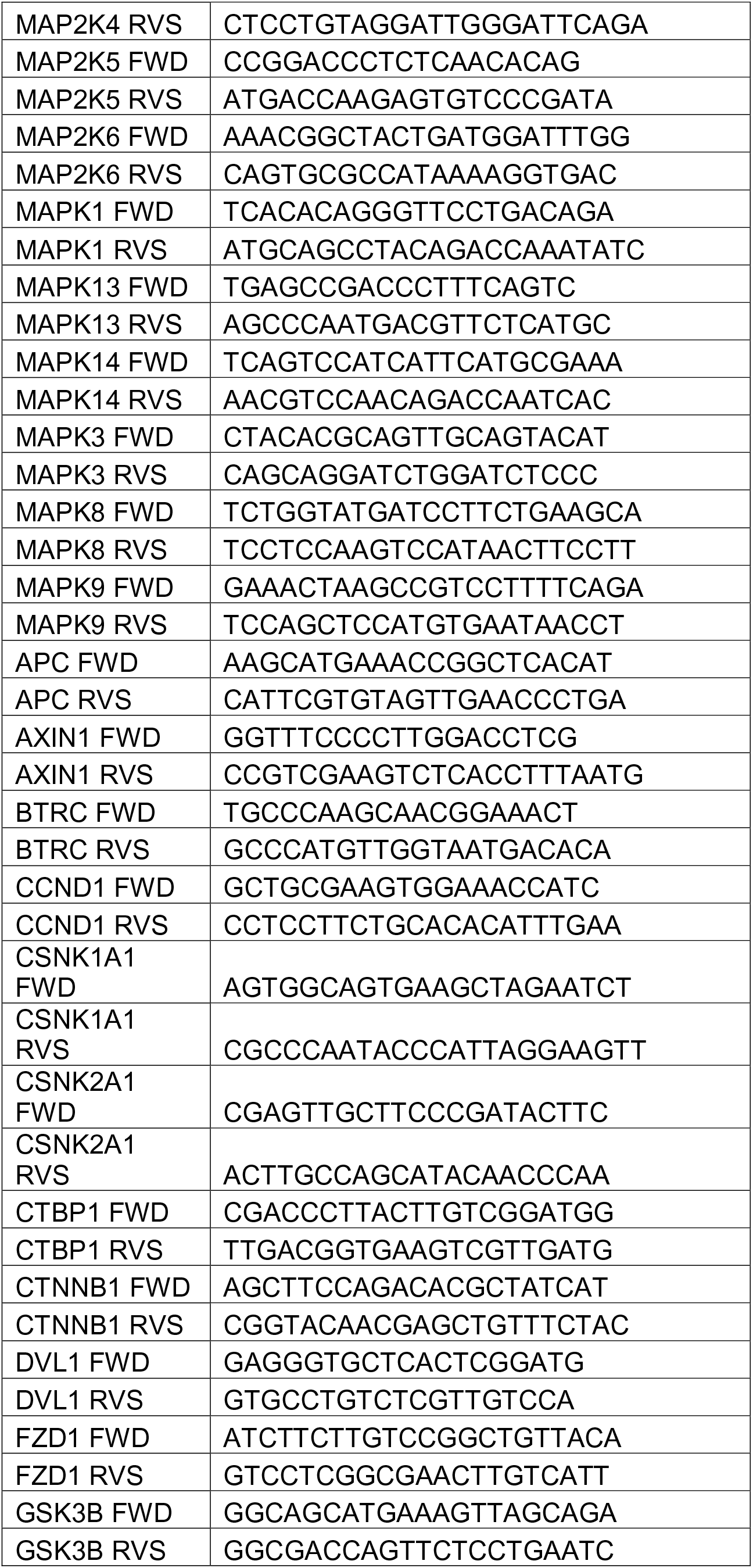
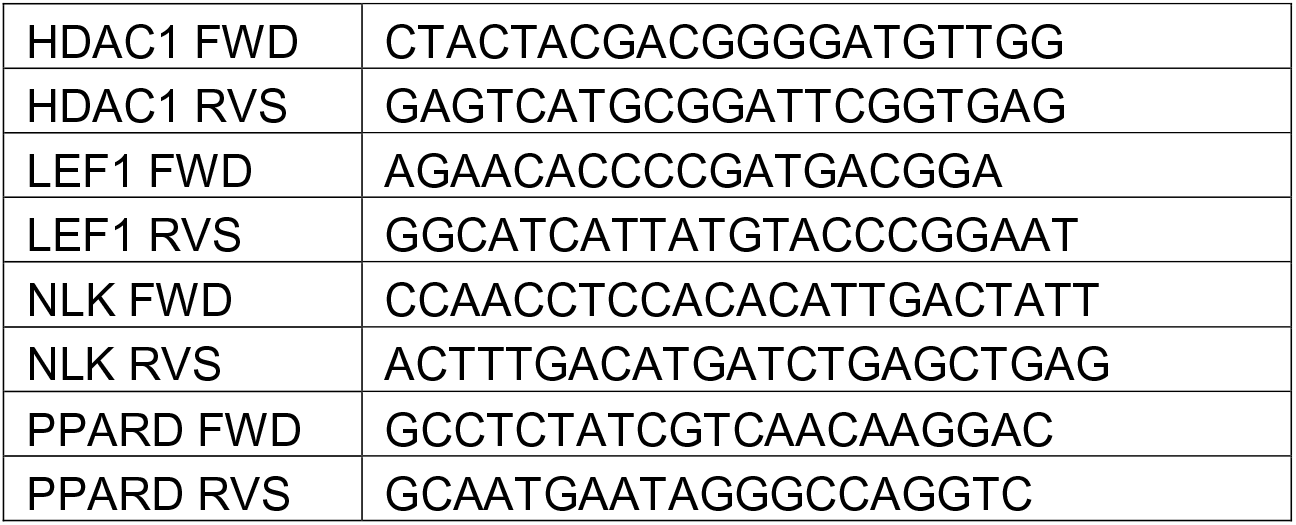
List of primers used to assess MAPK and WNT signaling pathways in the spheroid system.

To assess the functional consequences of MAPK and WNT pathway activation, we treated the tumor spheroids with nine small-molecule inhibitors that targeted key molecules of these pathways (Table 2). We quantified the volume change for each compartment over the course of the experiment, as assessing the inner core volume change from day 1 to day 7 is analogous to using the PrestoBlue viability assay^24^ to calculate fold increase for this spheroid model (Figure 2b, Extended Data Figure 2b).

**Table 2:** List of inhibitors in functional assessment of MAPK and WNT pathways in spheroid model

| Inhibitor | Target |
| --- | --- |
| SP600125 | JNK |
| BI-78D3 | JNK |
| PD0325901 | ERK |
| SB203580 | p38 |
| Vemurafenib | B-Raf |
| MK-2206 | AKT |
| IWR-1-endo | Axin2/β-catenin |
| LY317615 | PKC |
| CHIR-98014 | GSK3β |

The outer collagen layer volume change captures if the cells are actively degrading and re-organizing the matrix. Inhibition of JNK, ERK, and GSK3β significantly slowed the growth of spheroids (Figure 2c, Table 2). We also observed these inhibitors resulted in fewer cells in the collagen outer layer, leading to less volume change over the duration of the experiment (Extended Data Figure 2c).

We further explored the ERK cascade in the MAPK pathway by studying the impact MEK inhibition using highly selective MEK inhibitor PD0325901, as it was the most effective at stopping E-cad-mediated tumor spheroid growth (Figure 2a, c).

PD0325901 is a small-molecule inhibitor that prevents ERK activation by targeting MEK1/2 kinase activity^18^, which activates ERK by phosphorylating sites Y204 and Y187^25^. Selective small molecule inhibitors were chosen because shRNA knock-down of MEK or ERK would be fatal to the cells^27^, and we wanted to assess the potential of targeting this pathway in pre-clinical models of breast cancer. Results showed that PD0325901 was effective at reducing proliferation at a dose as low as 1 µM (Extended Data Figure 2d). E-cad+ spheroids featured increased levels of phosphorylated ERK (phospho-ERK) at sites Y187 and Y204 compared to E-cad- spheroids in all tested cell pairs (Figure 2d). We also observed a 3.5-fold increase in the IC_50_ value when E-cad was expressed (Figure 2e), indicating that more MEK inhibitor was required when E-cad was present to achieve the same decrease in cell growth. We confirmed this result using the more potent but less specific MEK1/2 inhibitor Trametinib^28, 29^ (Extended Data Figure 2e). We observed that MEK1/2 inhibition resulted in decreased phosphorylation of ERK but did not impact the phosphorylation of MEK (Figure 2f and Extended Data Figure 2f). Together these results suggest that E-cad expression induces a hyper- proliferative phenotype in breast cancer cells through ERK activation.

Next, we sought to understand if this activation of ERK relied on the well-known mechanical role of E-cad, that of promoting cell-cell adhesion. We treated E-cad+ cancer cells with the functional blocking antibody HECD-1 to prevent cells from making intercellular adherens junctions^30^. We first confirmed that the functional blocking antibody prevented cell-cell adhesion in E-cad+ cells via live cell microscopy over the incubation period: cells became more mesenchymal in morphology, similar to MDA-MB-231 E-cad- cells (Figure 2g, Extended Data Figure 2g). Also, as anticipated^22, 23^, in HECD-1-treated cells, actin filament organization was lost (Extended Data Figure 2h). Intriguingly, phospho-ERK expression measured via immunofluorescence revealed a shift from the cytoplasm to the nucleus after treatment with HECD-1 (Figure 2h). We also observed a decrease in the areas of the nucleus and the cytoplasm of HECD-1- treated cells (Extended Data Figure 2i).

### E-cad interacts with EGFR and induces hyper-activation of ERK

To further explore the mechanism by which E-cad expression leads to ERK activation and hyper-proliferation in breast cancer cells, we designed a bilateral orthotopic *in vivo* study. We injected E-cad+ and E-cad- cell pairs into the mammary fat pad of NOD-SCID Gamma (NSG) mice utilizing bilateral injection to allow for a direct comparison of the size of the two resulting tumors (Figure 3a). In all three tested cell pairs, we observed that E-cad+ tumors significantly grew more rapidly than E-cad- tumors, regardless of the level of endogenous E-cad expression of the parental cells (Figure 3b-d). Specifically, we observed a three-fold increase in the growth rate in E-cad+ KI MDA-MB-231 tumors (Figure 3b). For both MDA-MB-468 and MCF7 cell pairs, E-cad- tumors were already significantly smaller compared to E-cad+ control tumors three weeks after injection (Figure 3c, d). In the MDA-MB-231 bilateral study, the average weight of final E-cad+ tumors was ten-fold higher than the weight of E-cad- tumors (Figure 3b, Extended Data Figure 3a). This is remarkable considering that wild-type (endogenously E-cad-) MDA-MB-231 cells already grow rapidly *in vivo* and are widely used for that reason^20, 31^. Similar trends were observed for MDA-MB-468 and MCF7 cell pairs (Figure 3c, d, Extended Data Figure 3b, c). We repeated the MDA-MB-231 study with a single tumor per mouse to ensure there was no crosstalk between the two implanted tumors and confirmed the results of the bilateral study (Extended Data Figure 3d).

**Figure 3:**
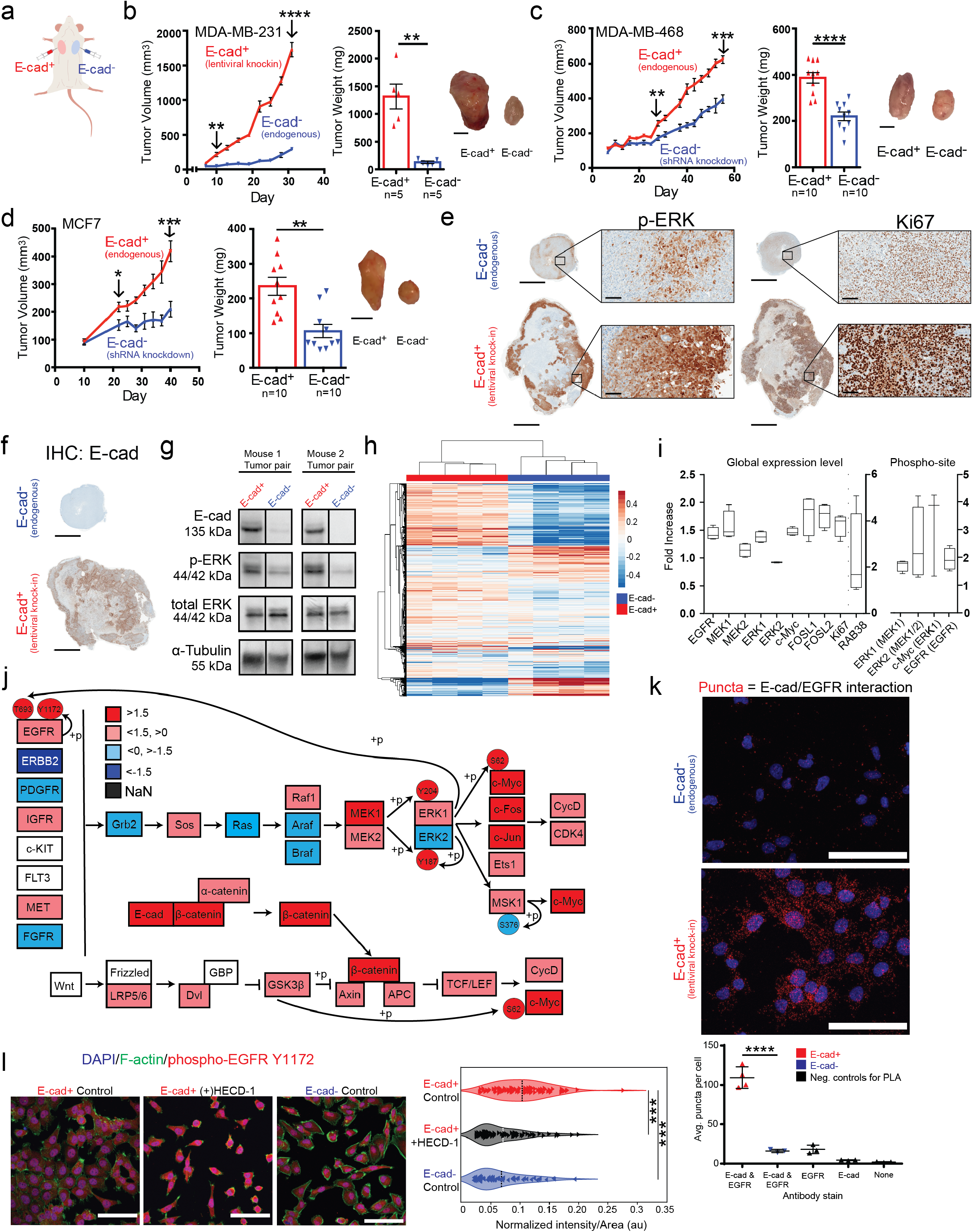
**E-cadherin expression correlates with increased proliferation and activation of MAPK pathway**. (a) Schematic of bilateral mammary fat pad injection for *in vivo* modeling of primary tumor growth. (b-d) Time-dependent tumor volume growth and final tumor weight of control MDA-MB-231 scramble and E-cad+ MDA-MB-231 tumors (****P< 0.0001) (b), control MDA-MB-468 scramble and E-cad- shRNA MDA- MB-468 tumors (***P= 0.0009) (c), and control MCF7 scramble and E-cad- shRNA MCF7 tumors (***P= 0.0003) (d). Apparent tumor volume was assessed using calipers to measure x and y dimensions. Tumor weight at the end of the study with pictures of representative excised tumors. Scale bars, 1 mm. **P=0.0079, ****P<0.0001, **P=0.0015 respectively. P values determined from paired T.Test for mice. (e) IHC staining of MDA-MB-231 bilateral tumor pair for phospho-ERK and proliferative marker Ki67 in MDA-MB-231 primary tumor pair. Scale bar = 3 mm, inset = 100 µm. (f) IHC staining for E-cad in MDA-MB-231 tumor pair. Scale bar = 3 mm, inset = 100 µm. (g) Western blot of two sets of primary tumors from the MDA-MB-231 bilateral mouse model, showing increased levels of phospho-ERK in E-cad+ tumors. (h) Heatmap of global site proteomics data comparing 4 MDA-MB-231 E-cad+ and E-cad- tumor pairs. (i) Box plots: fold change of key proteins in the MAPK pathway at the global proteomics level and activation differences of key proteins in the MAPK pathway (phosphosite proteomics data). (j) MAPK and WNT pathway schematics of fold change comparing MDA-MB-231 E-cad+ tumor to E-cad- tumor. Expression differences of +/- 1.5 fold change were considered biologically relevant. (k) Proximity ligation assay results demonstrate MDA-MB-231 E-cad+ cells have many E-cad/EGFR interactions, visualized by red puncta. Scale bar = 50 µm. Plot of average puncta/cell counts in E- cad+ and E-cad- cells, as well as in experimental negative controls (E-cad+ cells with single or no antibodies). N =2, n = 2 to 4. (l) Immunofluorescence staining of phospho- EGFR at site Y1172 in E-cad+ control and HECD-1 (functional blocking antibody) treated cells. Nuclear expression is unchanged, but cytoplasmic expression is decreased in HECD-1 treated E-cad+ cells. N =3, n = 3.

Immunohistochemistry (IHC) analysis of sections of MDA-MB-231 tumors from the bilateral study indicated that E-cad+ tumors displayed much higher levels of phospho-ERK and increased expression of the proliferation marker Ki67 compared to E- cad- tumors (Figure 3e). The regions of the tumor with high levels of expression of phospho-ERK and Ki67 correlated with the regions of the tumor with high levels of E- cad expression (Figure 3f). Furthermore, we performed western blotting analysis of the paired tumors from the bilateral study and found that the level of phospho-ERK was significantly higher in E-cad+ tumors compared to E-cad- tumors, with no significant change in total ERK (Figure 3g). This data suggests that E-cad expression impacts tumor growth by inducing cancer cell hyper-proliferation via ERK phosphorylation.

However, this may not be the only mechanism underlying hyper-proliferation in E-cad+ breast tumors. Hence, we conducted global and phosphosite proteomic analysis on MDA-MB-231 tumor pairs to assess all pathways activated by E-cad expression (Figure 3h). We designed this combined analysis to directly compare E-cad+ *vs*. E-cad- tumors obtained from the same mouse. A total of 7766 global proteins and 7258 phosphosites were identified. Using a cut-off of 1.5-fold relative expression change, we observed a ∼15% increase in global proteins expressed and ∼8% increase in activation sites in E-cad+ tumors compared to E-cad- tumors (Extended Data Figure 3e). We first use this data to confirm the MAPK pathway is activated by E-cad expression and map it more fully. Global proteomic analysis of the MAPK pathway revealed a 1.8-fold increase in the expression of oncogenic Ras protein (RAB38) and an increase in the expression of transcription factors downstream of ERK, including c-Myc (encoded by *MYC*) and FOS-related genes *FOSL1* and *FOSL2* (Figure 3i). RT-qPCR confirmed that FOS was also the most upregulated gene in the MAPK pathway at the mRNA level (Figure 2a). Phosphoproteomic analysis showed significant increases in the phosphorylation abundance of key activation sites in the MAPK pathway: Y1172 and T693 on EGFR, Y204 and Y187 on ERK1/2 and S62 on c-MYC (Figure 3i).

Next, we sought to determine if the MAPK and WNT pathways were the most changed when E-cad expression was manipulated. Utilizing the pathway analytical tools DAVID and KEGG^29–33^, we consistently found that the MAPK and WNT signaling pathways were the most highly upregulated among cancer-related pathways, which was one of the most upregulated pathway groups as determined by functional annotation (Figure 3j, Extended Data Figure 3f, g). This indicates that the MEK/ERK cascade is the driving force behind this hyper-proliferative phenotype induced by E-cad expression. The various protein and phosphosite changes around the ERK complex revealed by global and phosphosite proteomic analysis were confirmed via western blotting of spheroid samples (Extended Data Figure 3h, Extended Data Figure 2F).

With evidence that the MAPK pathway is the main driver of this hyper- proliferative phenotype associated with E-cad expression, we examined the cell membrane receptors that feed into the ERK cascade. Global and phosphosite proteomics revealed that the cell receptor EGFR was increased 1.4-fold in E-cad+ breast tumors compared to E-cad- tumors and its phosphosites Y1172 (via EGFR) and T693 (via ERK1/2) were upregulated 1.8-fold and 1.6 fold, respectively (Figure 3i, j). Phosphosite Y1172 is associated with EGFR autophosphorylation and phosphosite T693 is a result of feedback activation loop via ERK1/2 signaling^7, 17^. Hence, we hypothesized that E-cad-induced EGFR autophosphorylation was induced by crosstalk between EGFR and E-cad receptors at the cell membrane, increasing activation of the ERK cascade in MAPK and culminating in the hyper-proliferative phenotype of E-cad+ cancer cells. We note that E-cad has previously been shown to impact EGFR activation similarly to EGF-induced activation at phosphorylation site Y1172, albeit using *in vitro* assays and with normal mammary epithelial cells^7, 17^.

To assess the putative interaction between E-cad and EGFR receptors, we conducted a proximity ligation assay and quantified the number of cellular interactions between these two proteins via immunofluorescence. This assay revealed there were significant interactions between E-cad and EGFR in E-cad+ cancer cells, which had an average of 129 puncta compared to an average of < 20 puncta in E-cad- cancer cells (Figure 3k and Extended Data Figure 3i). We also assessed the protein level of EGFR in spheroids and observed significantly more EGFR in E-cad+ spheroids compared to E-cad- spheroids (Extended Data Figure 3h). We also stained control and HECD-1- treated E-cad+ cells with an antibody against phospho-EGFR Y1172 and demonstrated that functionally blocking E-cad disrupted EGFR activation, as there was less phospho-EGFR at the cell membrane in E-cad+ cells treated with HECD-1 (Figure 3l, Extended Data Figure 3j). Together these results show a functional relationship between E-cad and EGFR signaling, as these receptors are in close proximity to each other at the cell membrane, and disruption of E-cad intercellular junctions results in a decrease in EGFR activation at Y1172.

### E-cadherin enhances metastatic outgrowth

In addition to its role in primary tumor growth, proliferation may occur at metastatic sites, so we wanted to assess the role of E-cad in metastatic outgrowth. We utilized the chicken chorioallantonic membrane (CAM) assay, in which intravital microscopy can be used to track cells as they extravasate, colonize, and proliferate^33,34,35^ (Figure 4a). To fully control the impact E-cad expression has on the cellular phenotype, we generated MDA-MB-231 cells that only express E-cad after receiving an extracellular stimulus. This tunable E-cad-zsGreen cells gain E-cad expression when activated by the cell-permeant high-affinity ligand Shield-1^36^. We set up three experimental conditions: (1) cells that did not express E-cad (E-cad- cells and cells expressing tunable E-cad treated with vehicle control DMSO), (2) cells that expressed E-cad at all times (E-cad+ cells), and (3) cells that did not express E-cad until after extravasation into the embryo (tunable cell line treated with Shield-1).

**Figure 4:**
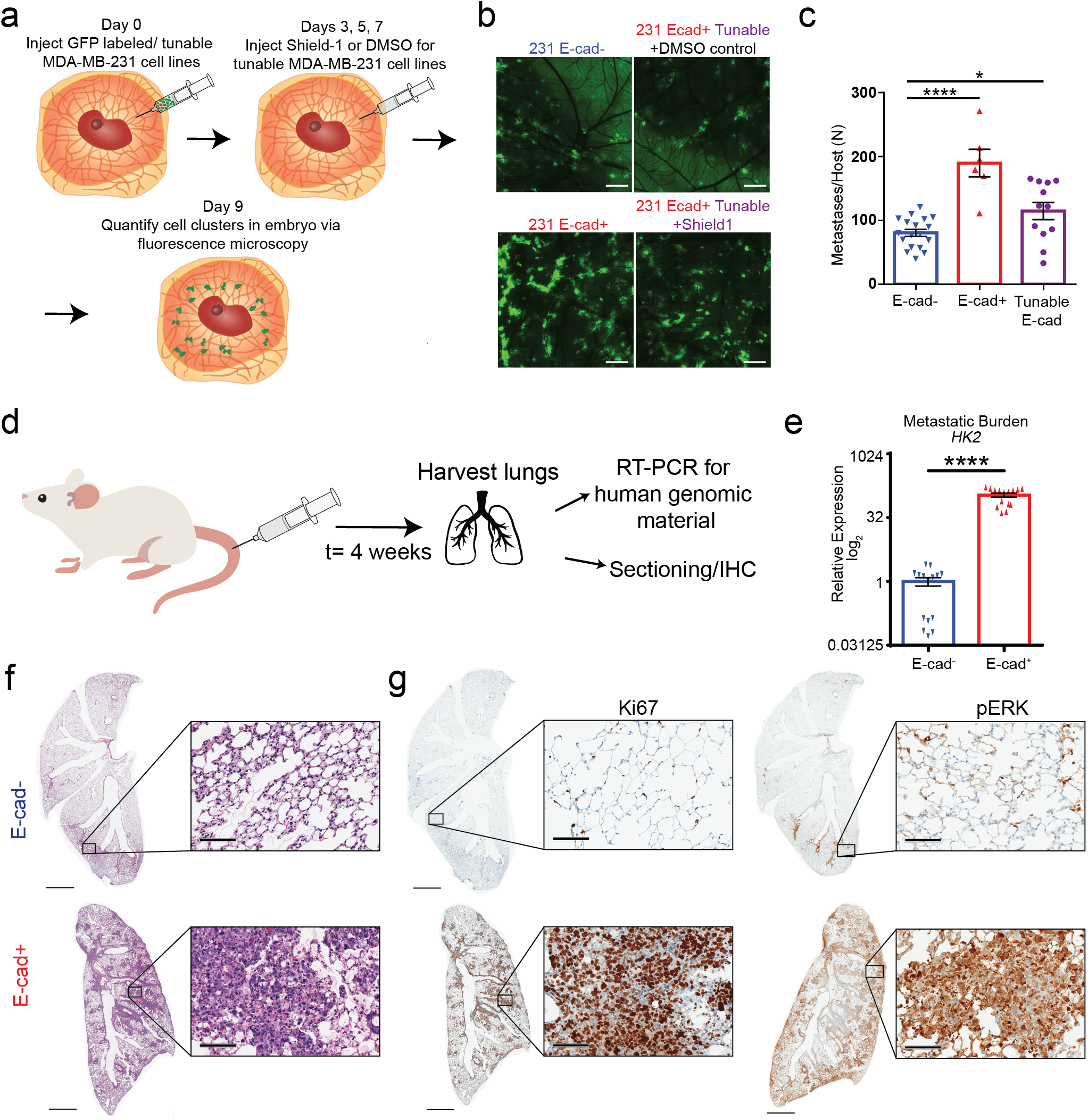
E-cadherin results in larger metastatic burden in two *in vivo* models via MAPK proliferation pathway. (a) Schematic of intravital microscopy experiment using CAM assay. (b) Confocal images of embryos on day 18 of embryo growth (9 days post injection) showing greater accumulation of GFP tagged E-cad+ and Shield-1 treated tunable cells when compared to the scramble control (E-cad-) or DMSO treated tunable cells. ROI 6x6 tile, 10x objective. (c) Number of metastases per host quantified from fluorescent images. (d) Schematic of tail vein injection of breast cancer cells. (e) Relative expression level of *HK2* a marker of human genomic material comparing lungs of mice injected with E-cad+ to E-cad- cells. (f) H&E staining of lungs from E-cad+ and E-cad- mice. Scale bar = 2 mm. (g) IHC staining for Ki67 and phospho-ERK (respectively) in the lungs of from mice injected with E-cad+ and E-cad- MDA-MB-231 cells. Scale bar = 2 mm.

To ensure that E-cad did not impact the colonization potential of cancer cells, we conducted an extravasation assay in the CAM system (Extended Data Figure 4a, b). Fluorescent images of CAM showed larger colonies in both embryos injected with E- cad+ cells and embryos injected with the tunable cells treated with Shield-1, when compared to E-cad- cells and embryos treated with DMSO (Figure 4b). When quantified, the E-cad+ conditions had a 1.5 to 2-fold increase in the number of metastases present throughout the CAM when compared to those lacking E-cad expression (Figure 4c). This data confirmed that E-cad expression impacted cancer cell growth after completion of the metastatic cascade (i.e., post-extravasation from vessels into embryo of CAM).

We also designed a more biologically relevant metastatic assay, the mouse tail vein injection, in which cancer cells were “forced” to colonize the lung via tail vein injection (Figure 4d). The lung is the most common site of metastasis for breast cancer in mice and is one of the most common sites in humans^31, 37^. We injected the same number of either E-cad+ or E-cad- breast cancer cells into the tail vein of NSG mice and assessed metastatic burden in the lung four weeks after injection. We chose an extended timeline to minimize effects of survival or colonization ability and focus on metastatic growth. Mice weights were tracked throughout the experiment to monitor health (Extended Data Figure 4c). After four weeks, we observed significantly more macro metastases in the lungs of mice injected with E-cad+ cells (Extended Data Figure 4d). RT-qPCR analysis of the human genomic marker *HK2*^32^ showed a remarkable 109-fold increase in the relative expression in lungs of mice injected with E-cad+ cells compared to Ecad- cells (Figure 4e). These results were confirmed via H&E staining of lung sections, where dense foci of cancer cells were present in the lungs of mice injected with E-cad+ cells (Figure 4f). In contrast to E-cad- cells, we also observed that a majority of the cells in the lungs of mice injected with E-cad+ cells were positive for Ki67 (Figure 4g). Finally, we observed a significant increase in phospho-ERK expression when E-cad+ cells were used for the experimental metastasis assay and maintained E-cad expression (Figure 4g, Extended Data Figure 4e). These results suggest that, in addition to its role in primary tumor growth, E-cad promotes proliferation at the secondary metastatic site.

### Blocking ERK phosphorylation reverses E-cad-induced cancer cell hyper- proliferation

To validate the above mechanism of action of E-cad and as a way to begin to test potential translation in the clinic, we subjected MDA-MB-231 E-cad+ tumor-bearing mice to MEK1/2 inhibitor PD0325901, selected from our *in vitro* drug screen (Figure 5a). We used PD0325901 also because it is frequently used when studying ERK activity due to its specificity to MEK1/2 kinase activity without impacting MEK activation^18^. This MEK inhibitor is administered orally to patients^38, 39^ and the dosage and scheme were determined based on clinical trials for solid or breast tumors and preclinical *in vivo* models^40^. Mice were given 20mg/kg of PD0325901 for 5 days and then had 3 weeks off before the next treatment was given to ensure safety of the animals given the clinical evidence of severe toxicity at high dose MEK inhibitor^41, 42^. We monitored the weight of the animals throughout the study, as it would be the first sign of toxicity and did not observe a significant impact on mouse weight with the 5-day administration (Extended Data Figure 5c).

**Figure 5:**
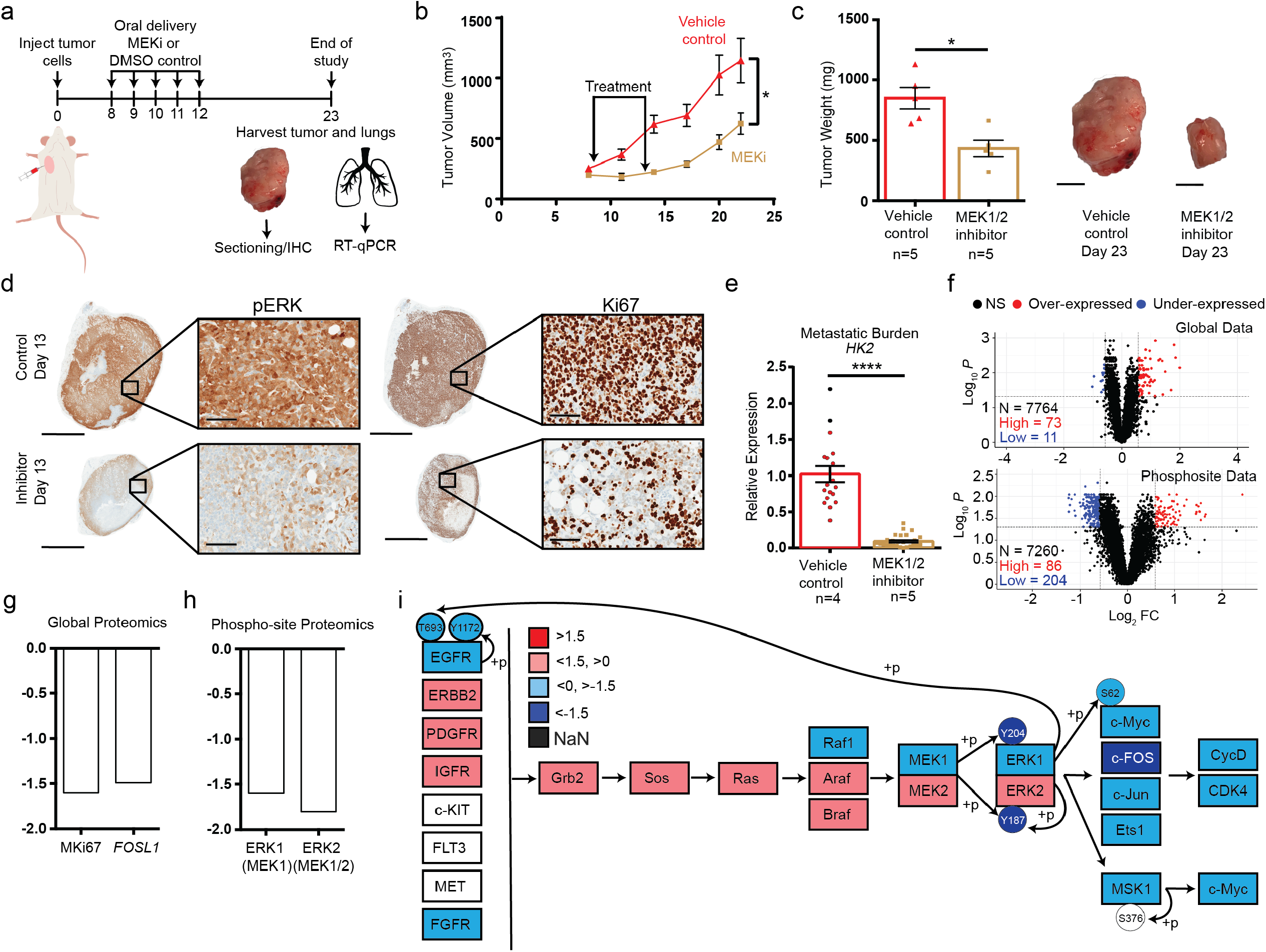
Blocking ERK phosphorylation stops hyperproliferative effect of E-cad. (a) Schematic of orthotopic injection into mammary fat pad of NSG mice to assess impact of MEK1/2 inhibitor on primary tumor progression (N=5 per group). (b) Time-dependent volume of MDA-MB-231 E-cad+ untreated (red line) and treated (gold line). Apparent tumor volume was assessed using calipers to measure x and y dimensions. *P =0.0345. (c) Tumor weight at the end of the study with representative pictures of excised tumors. *P=0.0159. Scale bar =1 mm. (d) IHC staining of tumors harvested on day 13, 1 day after treatment was completed, showing downregulation of phospho-ERK with treatment of MEK1/2 inhibitor PD0325901. There is a significant decrease in Ki67 positive cells when mice were treated with MEK1/2 inhibitor PD0325901. Scale bar = 3 mm. Inset = 100 µm. (e) Relative expression level of *HK2* (marker for human genomic material) in the vehicle control and MEK1/2 inhibitor treated groups. ****P<0.0001. N=2, n=3 per mouse for qPCR. (f) Volcano plots showing statistically significant fold increase/decrease during treatment in the global proteomics and phosphosite proteomics. (g) Fold decrease in global protein levels of proliferation marker Ki67 (P= 0.0001) and FOS related protein encoded by FOSL1 (P= 0.0015). (h) Fold decrease in phosphosites Y187 (P= 0.254) and Y204 (P= 0.053) on ERK1/2 when treated with MEK1/2 inhibitor P0D0325901. (i) Pathway schematic comparing relative expression of treated tumors to control tumors.

Compared to vehicle control, we observed a significant decrease in tumor progression and final tumor weight and size when mice bearing E-cad+ tumors were treated with PD0325901^11^ (Figure 5b, c). After treatment ended on day 12, the growth curve of the treated tumors increased again, as anticipated. We harvested tumors on day 13, the day after treatment ended, and observed the anticipated decrease in phospho-ERK and Ki67 levels via IHC staining (Figure 5d). Note that for tumors harvested at the conclusion of the study (day 23), phospho-ERK and Ki67 levels were not significantly different (Extended Data Figure 5b), further explaining the increase in tumor growth after treatment concluded on day 12. Hence, MEK inhibition via PD0325901, which prevents ERK activation, can slow E-cad+-induced tumor hyper- growth (Extended Data Figure 5d).

We also assessed metastatic burden in the lungs using RT-qPCR and observed a 90-fold decrease in relative expression of *HK2*^32^ when mice were treated with the MEK1/2 inhibitor compared to control mice (Figure 5e). Hence, MEK1/2 inhibition slows down E-cad-induced growth at both the primary and secondary sites. These results also suggest that E-cad+ cells employ the same proliferative pathway at metastatic sites, as observed in mice lungs stained for phospho-ERK (Figure 4g.

To ensure that the inhibitor PD0325901 worked specifically and did not significantly impact other pathways, we conducted global and phosphosite proteomics on the treated and untreated tumors. We observed that the expression of only ∼1% of proteins and ∼4% of phosphosites were impacted by PD0325901 treatment when comparing treated to control E-cad+ tumors (Figure 5f). The expression of FOS-related gene *FOSL1* and Ki67 decreased in the treated mice when compared to control mice (Figure 5g). Analysis of the phosphosite data confirmed that the inhibitor prevented phosphorylation of ERK by MEK1/2 at sites Y187 and Y204 (Figure 5h). We also observed a slight decrease in EGFR and phosphosite Y1172 in treated tumors (Figure 5i). After assessing the impact of treatment on pathway dynamics, we observed that the inhibitor worked as intended and specifically, decreasing MEK activation of ERK at sites Y204 and Y187 (Figure 5i). These results confirm the results obtained using the 3D spheroid model: E-cad+ spheroids treated with PD0325901 featured decreased levels of phospho-ERK activation at sites Y204 and Y187 (Extended Data Figure 2f). We also conducted western blot analysis for EGFR levels in spheroids after treatment with PD0325901 and observed a decrease in EGFR protein in treated spheroids (Extended Data Figure 3h). This data demonstrates the pathway is preserved *in vivo*, in addition to the response to treatment.

## Discussion

E-cadherin is an EMT marker whose loss is correlated with successful cancer metastasis^1, 2, 22, 23^. This characterization has led to its classification as a tumor suppressor gene. In breast cancer, this mechanistic understanding seems to contradict clinical data, however, as patients with E-cad positive tumors have reduced survival rates^5^. This discrepancy has provoked a reassessment of the role of E-cad in cancer, mostly with a focus on the protein’s role in EMT and metastatic disease^11–14^. Based on its interactions in normal mammary epithelial cells with receptor tyrosine kinases (RTKs), which feed into various cell growth and proliferation pathways, here we assessed the role of E-cad in tumor progression as a driver of cancer-cell proliferation^6^. Through our recently developed two-compartment tumor spheroid model and *in vivo* models, we identified a previously unrecognized hyper-proliferative cascade that critically depends on E-cad expression and may be responsible for the highly aggressive nature of E-cad positive tumors.

Due to its mechanical role as a cell-cell adhesion molecule, it was imperative to study E-cad in a biologically relevant environment, ensuring E-cad can create adherens junctions in all directions rather than be limited to planar interactions in standard culture conditions. Utilizing a 3D double layered spheroid system, we observed large proliferation changes in our engineered cell pairs. In animal models, we observed that E-cad expression correlated not only with increased primary tumor growth, but also with metastatic outgrowth. This data suggests that although E-cad has implications in the migratory and invasive phenotypes of cancer cells, the impact E-cad expression has on proliferation is the larger driving force in breast cancer progression.

Our phosphoproteomic assessment identified ERK1/2 activation through the MAPK signaling cascade as the molecular mechanism contributing the most to this dramatic change in proliferation. This is supported by recent studies which show that, in normal mammary epithelial cells, ligated E-cad can recruit EGFR, induce its ligand-independent activation, and result in activation of the MAPK pathway^27, 31^. We identified MEK1/2 phosphorylation of ERK at Y204 and Y187, and auto-phosphorylation of EGFR at Y1172 as the key E-cad-dependent activation sites in the MAPK pathway. This phosphosite on EGFR is activated in the presence of EGF or similar stimulus, like soluble E-cad^10^, in mammary epithelial cells and has been linked to autophosphorylation behavior of EGFR at site Y1172^17^. This phosphosite can feed into either the MAPK or PI3K pathway, resulting in changes in cell cycle, growth, proliferation, metabolism, etc.^6, 7, 10, 17^ Our proteomics assessment allowed us to mechanistically connect these disparate observations and demonstrate that the ERK cascade in MAPK pathway is the most important driver behind the hyper-proliferative phenotype we observed in E-cad+ breast cancer cells. Functional blocking of the E-cad–EGFR interaction changed the localization of phospho-ERK to be more abundant in the nucleus as well as resulted in a decrease in phospho-EGFR (site Y1172) localized at the cell membrane. These results indicate E-cad must be functional for EGFR to be stimulated and undergo auto- phosphorylation; thus, providing further evidence that supports functional crosstalk between cadherins and RTKs exists in cancer^6, 17^. These results also provide evidence of the downstream impacts these cell surface interactions have on tumor progression – causing breast tumors to enter a state of hyper-proliferation^6, 17^.

Pharmacological inhibition of MEK1/2 kinase activity with PD0325901 inactivated ERK1/2 and is sufficient to reverse this hyperproliferative effect *in vitro* and *in vivo*. This inhibitor reverted E-cad+ primary tumor growth to that of E-cad- tumors and decreased metastatic burden in the lungs. Using inhibitors that specifically target the MEK1/2 kinase activity hold potential as a treatment option for E-cad positive breast tumors; although, delivery is an obstacle due to the side effects and off target impacts of MEK inhibitors on other organs and areas of the body. Clinical trials for PD0325901, previously tested for efficacy in treating solid tumors of the breast and melanoma, had to be terminated due to toxicity concerns and severe side effects^38–42^, Nevertheless, MEK inhibitors are only approved for combination treatment of a specific subset of melanoma. Our work suggests that patients with E-cad+ breast tumors could benefit from treatment with MEK inhibitors.

Clinical data examining IDC, which accounts for 80% of all breast cancers, and of which 90% are E-cad+, corroborates our results: patients with high E-cad expression have a worse overall survival^5^. It was suspected that there was a role between E-cad and MEK/ERK activation in other cancer types when a similar phenomenon was observed in the SKOV3 cell line. ^30^ Here we identify a molecular mechanism responsible for these observations and a therapeutic target with implications that begin but do not end with breast cancer.

---

**Extended Data** **Fig 1**: (a) Endogenous expression of E-cadherin in commonly used breast cancer cell lines. (b) 2D cell proliferation assay utilizing PrestoBlue to calculate the fold increase of the cells from day 7 to day 1. N = 3 biological repeats. *p =0.01, **p = 0.003, ***p = 0.0002.(c) Standard curves for utilizing PrestoBlue as a way to measure fold increase of spheroids. Shows strong correlation of signal with seeding density in the spheroid cores. (d) DIC confocal images of spheroid growth in all engineered cell lines at various time points. Images taken at 10x and 20x for zoom windows. Scale bar represents 200 µm and zoom window scale bar represents 50 µm. All data are mean ± SEM. P values determined Mann-Whitney test, two sided. (e) Ki67 and cleaved-caspase 3 staining of MDA-MB-231 E-cad- spheroid on day 5 of culture. Split channel of MDA-MB-231 E-cad+ spheroid on day 5 stained for Ki67 and cleaved-caspase 3, shown in figure 1. Scale bars: 200 µm, inset: 50 µm.

**Extended Data** **Figure 2**: (a) Relative expression of genes in the MAPK and WNT signaling pathways in MDA-MB-231 E-cad+ to MDA-MB-231 E-cad- cells (qPCR). Red bars represent 2- fold increase or greater, blue bars represent 2-fold decrease or less, and black bars represent value in between +2 and -2-fold increase. We considered a fold increase/decrease of higher/less than 2 to be biologically relevant. (b) Resulting growth curves of MDA-MB-231 E-cad+ and E-cad- cell line pair assessed via PrestoBlue and volume tracking. (c) Outer volume ratio of 10 µM inhibitor screening (N=1, n=5-6 per condition). (d) PD0325901 treated spheroids inner volume ratio with 1 µM and 10 µM inhibitor screening (N=1, n= 3-5 per condition). PD0325901 treated spheroids outer volume ration with 1 µM and 10 µM inhibitor screening (N=1, n=3-5 per condition). (e) IC_50_ of Trametinib E-cad+ cells require 3x more inhibitor to achieve the same effect of MI4 in E-cad- cells. p value <0.0001. (f) Western blot of MDA-MB-231 spheroids treated with MEK inhibitor PD0325901, demonstrating specificity to ERK inhibition, as phospho-MEK levels are unchanged in treated E-cad+ spheroids. (g) Images of MDA-MB-231 E-cad- cell migration at t = 2 min, t = 2 h, and t = 4 h. Negative control for images shown in main figure. Scale bar = 50 µm. (h) Split channel immunofluorescence of MDA-MB-231 E-cad+ cells shown in main figure for phosphorylated ERK (red), shown with nuclear stain (blue) and f-actin stain (green). Scale bars = 50 µm. (i) Area distribution of nucleus and cytoplasm in the 2 conditions indicating a decrease in both areas in the presence of E-cad functional blocking antibody.

**Extended Data** **Figure 3**: (a) Excised tumors from MDA-MB-231 bilateral study (4 out of 5 tumors). (b) Excised tumors from MDA-MB-468 study (7 out of 10 tumors). (c) Excised tumors from MCF7 study (10 out of 10 tumors). (d) Single tumor growth curves of MDA-MB-231 E-cad- and E-cad+ cell lines in NSG mice and final tumor wright to compare to bilateral tumor study results. (e) Volcano plots of global and phosphosite proteomics data, demonstrating the number of proteins with biologically relevant (+/- 1.5 fold change) when E-cad expression is manipulated in MDA-MB-231. (f) Heatmap of the top proliferation pathways in cancer as defined by KEGG database, demonstrating MAPK is one of the most upregulated pathways in E-cad+ breast tumors. (g) DAVID bioinformatics output of the upregulated genes in E-cad+ tumors as determined via global proteomics for functional annotation chart utilizing KEGG pathways. (h) Western blot assessment of MDA-MB-231 spheroids treated with PD0325901 demonstrating EGFR expression is impacted when ERK activity is inhibited. (i) Proximity ligation assay full panel of negative controls: MDA-MB-231 E-cad+ cells stained with each antibody individually and with no antibodies as well as computational image analysis quantification of puncta per cell. Each symbol represents the average puncta/cell for 1 well. Each well had approximately 30 cells analyzed. Scale bars = 50 µm. (j) Split channel of actin and phospho-EGFR Y1172 immunofluorescence stain in E-cad+ control and HECD-1 treated cells. Scale bars = 50 µm.

**Extended Data** **Fig 4**: (a) Schematic of CAM extravasation assay. (b) Extravasation rate of E-cad-, E-cad+, and tunable E-cad MDA-MB-231 cells. (c) Weights of mice for duration of the tail vein study. (d) Excised lungs showing visible nodules. (e) IHC of excised lungs, stained for E-cadherin. Scale bars are 2mm, zoom scale bars are 100 µm.

**Extended Data** **Fig 5**: (a) Pictures of excised tumors at the conclusion of the study (day 23). (b) IHC staining for phospho-ERK and Ki67 of tumors harvested on day 23. (c) Mouse weights throughout the study, showing weight loss in treatment group during MEK inhibitor administration (day 8-13). (d) Tumor volume progression of MDA-MB-231 E-cad+ compared to MDA-MB-231 E-cad- treated with PD0325901, demonstrating effectiveness of MEK inhibitor PD0325901 in reversing the hyperproliferative phenotype induced by E-cad expression.

## Materials and Methods

### Cell culture

Human breast carcinoma cells MDA-MB-231, MCF7, MDA-MB-468, (ATCC) were cultured in Dulbecco’s modified Eagle’s medium (DMEM, Corning, 10-013-CV) supplemented with 10% (v/v) fetal bovine serum (FBS, Corning, 35-010-CV) and 1% Penicillin-Streptomycin (Gibco, 15140-122). Cells were maintained at 37°C and 5% CO_2_ in a humidified incubator during cell culture.

### Lentivirus production and transduction

HEK-293T cells (ATCC) were cultured in Dulbecco’s modified Eagle’s medium (DMEM, Sigma Aldrich) supplemented with 10% (v/v) fetal bovine serum (FBS, Corning) and 1% Penicillin-Streptomycin (Corning). Mission shRNA glycerol stock for CDH1 (NM004360, TRCN0000237841) was obtained from Sigma. Bacterial culture was started and incubated for 16 h (overnight) on a shaker at 37°C and a speed of 200 RPM. Plasmid DNA was extracted using Midi-Prep Kit from (Machery-Nagel, 740412.50). Calcium chloride (2M) and HEPES-buffered-saline were used for transfection and lentiviral production. Plasmid DNA and reagents were added to HEK-293T cells and virus was harvested every 24 hours for 3 days. E-cadherin positive cells (MCF7, MDA-MB-468) received two rounds of virus for a total period of 48 h. After 48 h, cells were given fresh DMEM. After 24-48 h, cells were given DMEM with 5µg/mL puromycin for selection. Knockdown cells were cultured in puromycin conditioned medium for three days before checking protein expression. Cells were kept in selection medium for the duration of the experiments. Lentiviral knock-in of MDA-MB-231 (endogenously no E-cad expression) was generated as previously described^1^. Briefly, the lentiviral vector of E-cadherin-EGFP was generated from EGFP; pCS-CG (Addgene), via cloning full-length E-cadherin upstream of EGFP between the Nhe1 and Age 1 sites of pCS-CG to generate an EGFP-fused protein^1^.

### 2D immunofluorescence staining and imaging

Cells were seeded at desired density in 96 well flat-bottom plate. After 24 h growth, 4% paraformaldehyde (Sigma) was added to each well to fix cells and removed after 20 min incubation at RT. After washes with DPBS, 0.1% Triton (Sigma) was added to cells to permeabilize the cell membrane and incubated for 5 min at room temperature before removal. Wells were then blocked with 5% Normal Goat Serum (NGS) (Sigma) for 1 h. Primary antibodies were added, and samples were incubated overnight in a 2-8°C fridge. After removal of the primary antibodies (see table 3), samples were stained with secondary antibodies in 1% NGS for 30 min at room temperature. Samples were then stained with H33342 (nuclear DNA) and phalloidin (F-actin) in 1% NGS for 15 min at RT. Samples were then washed 3x with DBPS (Corning) before imaging. Confocal images were captured using a 40x oil objective and Nikon A1 Confocal microscope. Immunofluorescence analysis was conducted by a CellProfilerTM pipeline. Briefly, nuclei and their corresponding cell bodies were masked from the DAPI and Phalloidin Channels respectively. These masks were applied to the Biomarker channel to elicit spatial expression at single-cell resolution.

**Table 3:**
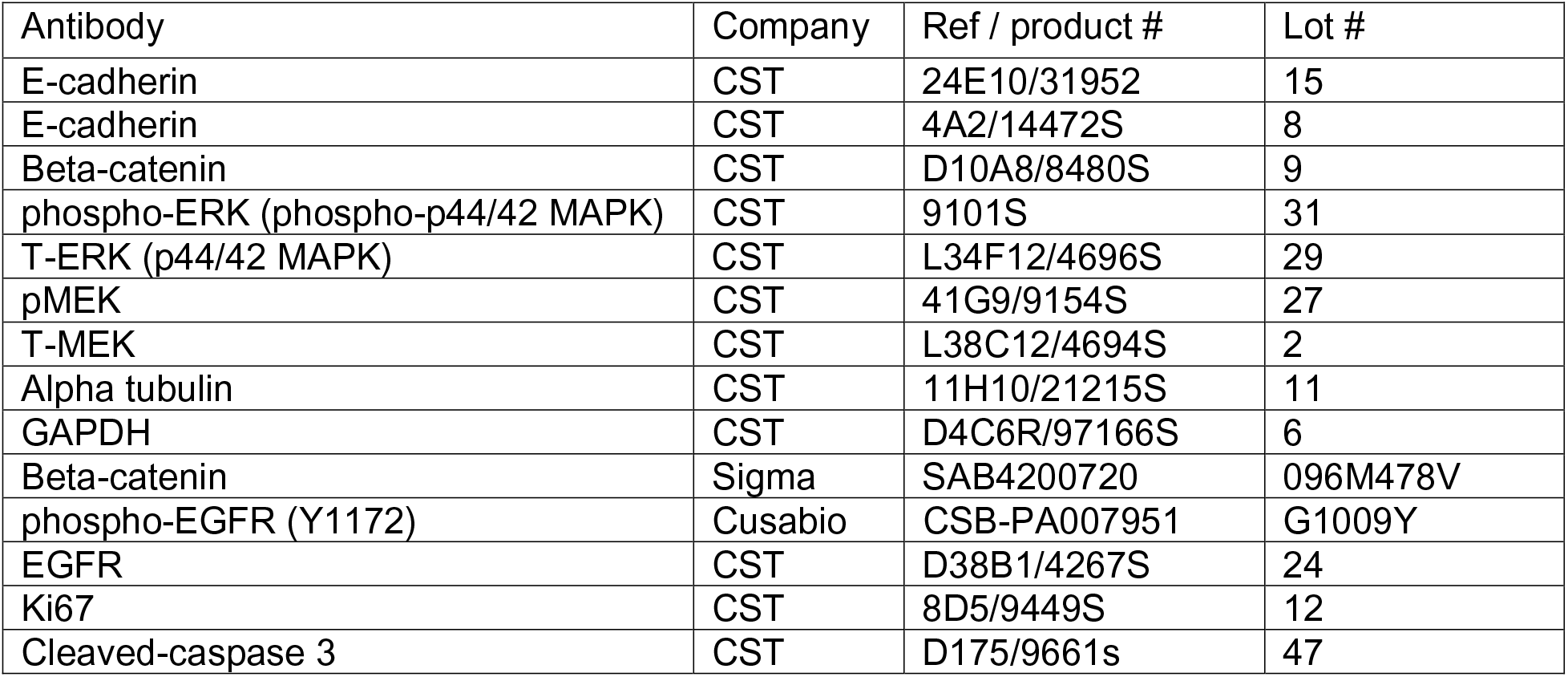
List of antibodies used in immunofluorescence, immunohistochemistry, and western blots

| Antibody | Company | Ref / product # | Lot # |
| --- | --- | --- | --- |
| E-cadherin | CST | 24E10/31952 | 15 |
| E-cadherin | CST | 4A2/14472S | 8 |
| Beta-catenin | CST | D10A8/8480S | 9 |
| phospho-ERK (phospho-p44/42 MAPK) | CST | 9101S | 31 |
| T-ERK (p44/42 MAPK) | CST | L34F12/4696S | 29 |
| pMEK | CST | 41G9/9154S | 27 |
| T-MEK | CST | L38C12/4694S | 2 |
| Alpha tubulin | CST | 11H10/21215S | 11 |
| GAPDH | CST | D4C6R/97166S | 6 |
| Beta-catenin | Sigma | SAB4200720 | 096M478V |
| phospho-EGFR (Y1172) | Cusabio | CSB-PA007951 | G1009Y |
| EGFR | CST | D38B1/4267S | 24 |
| Ki67 | CST | 8D5/9449S | 12 |
| Cleaved-caspase 3 | CST | D175/9661s | 47 |

**Table 4:**
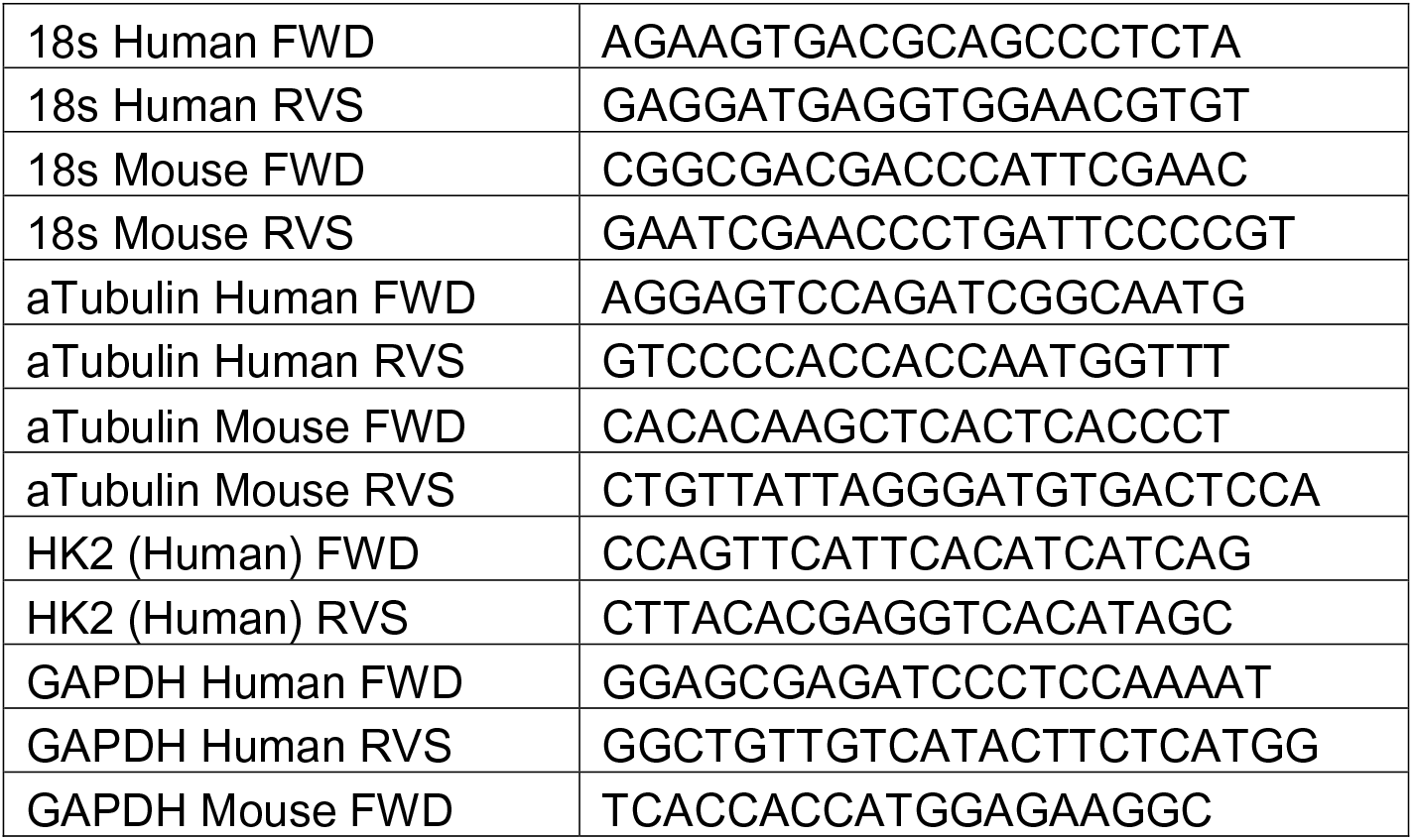
List of primers used to assess metastatic burden (*HK2*).

### Spheroid modeling

A double-layered spheroid model was generated with Matrigel (Corning) and rat tail type I (Corning). Cells were trypsinized, pelleted, and re-suspended in Matrigel at the desired cell density and then wrapped in a collagen I gel was mixed as previously described by Fraley *et al* ^2^. The spheroids are plated in a 96-well round-bottom dish and suspended in 200 µL medium.

### Spheroid immunofluorescence staining and imaging

Spheroids were harvested at various time points, washed 3x in DPBS (Corning) and then fixed in 4% paraformaldehyde (Sigma) overnight at 4°C. Spheroids were rinsed with DPBS prior to capturing DIC images using Nikon A1R confocal microscope. Spheroids to be stained with antibodies were permeabilized with 0.1% Triton-X (Sigma) for 10 min. Spheroids were then washed with DPBS and blocked with 5% Normal Goat Serum (NGS) (Cell Signaling Technology) for 1 h. Primary antibodies were added at 1:100 dilution and incubated overnight at 4°C. Following primary antibody incubation, spheroids are washed 3x in DPBS and then stained with secondary-antibodies (Invitrogen) as well as F-actin/phalloidin (Invitrogen) and Hoechst 3342 (ThermoFisher) in 1% NGS for 1 h at RT. After fluorescent stain incubation, spheroids are washed 3x with DPBS and then incubated in tissue clearing solution (60% glycerol, 2.5 M fructose in DI water) for 20 min at RT. Wash spheroids 3x before imaging on Nikon AX confocal microscope using 20x water immersion objective and 40x water immersion objective.

### 2D and 3D Proliferation using PrestoBlue viability assay

Cells were seeded in 96 well plate at 2,000 cells/well for MDA-MB-231 and MCF7 and at 5,000 cells/well for MDA-MB-468 (day 0). On day 1 and day 7, cells were incubated in 1x PrestoBlue (ThermoFisher) at 37°C for 1 h. To account for background signal, 5 to 10 wells of 1x PrestoBlue were prepared and measured to subtract off background medium signal from the experimental wells. After reading the signal, cells were washed with DPBS 3x, given 100 µL of fresh medium.

Spheroids were incubated at 37°C in a 1x PrestoBlue for 3 hours on day 1 (day of generation). Standard curves were generated relating cell number to RFU to ensure linear relationship (Extended Data Figure 1 e-g). After reading, spheroids were wash in DBPS 5x and given 200 µL of fresh medium. Experiment was carried out for either 5 or 8 days. To account for background signal, 5-10 wells of 1x PrestoBlue were prepared and measured. The average background signal value was subtracted from the experimental wells. SpectraMax plate reader (Molecular Devices) was used to read the RFU for all experiments at (EX 540, EM 600, cutoff 590).

### mRNA extraction and cDNA synthesis

Spheroids were generated and then incubated in fresh media. After 24 h, spheroids were harvested and washed 3x in DPBS (Corning). To isolate mRNA, TriZol reagent (Invitrogen) was added to the washed spheroids and then vortexed until completely dissolved. Direct-zol Mini Prep kit (Zymo research) reagents and protocol were used to complete RNA extraction. cDNA was synthesized using the Bio-Rad cDNA iScript kit following the recommended protocol to make 1000 ng of cDNA for a total reaction volume of 20 µL.

### RT-qPCR

qPCR was conducted with iTaq-SYBR Green (Bio-Rad) using the Bio-Rad CFX CFX384 Touch Real-Time PCR detection system. Primers were obtained from Integrated DNA Technologies and are listed in Table 1.

### Inhibitor studies

Inhibitor studies were conducted using spheroids described above. Experiments were carried out for a week and imaged daily using phase contrast microscopy. Inhibitors were obtained from Selleckchem (Table 2) and fresh or conditioned medium (containing drug) was added on the day the spheroids were generated and not changed during the week-long experiment.

### Quantifying double-layered spheroid volume

Inner and outer layers of the spheroids were manually traced using NIS Elements software to generate a radius measurement. Volume of inner and total spheroid was calculated using the volume for a sphere: V=4/3 πr . To calculate the volume of the collagen layer, the inner volume was subtracted from the total volume.

### Protein extraction, concentration, and western blot assays

A clear sample buffer containing SDS with protease inhibitor (Roche, 05892970001) was used to lyse 10-15 spheroids or 2D cells via cell scraping method from a confluent 10cm dish. Samples were sonicated for 10 -15 pulses using a needle probe sonicator (VWR Scientific). Samples were then heated at 100°C for 5 min to completely denature the protein. Protein samples were stored in -80°C freezer until ready for use. Protein concentration was measured using MicroBCA Kit (ThermoFisher, 23235) following the manufacturer’s protocol. A loading buffer made from 2x Laemlli (Biorad) was prepared with the instructed amount of beta-mercaptoethanol and added to the protein samples for a final concentration of 2 µg/µL. Bio-rad mini-protean gels (4-15% gradient, 4561086) were used for gel electrophoresis. Transblot Turbo packs (Bio-rad) were used to transfer the protein from the gel to PVDF membrane. Antibodies are listed in Table 3 and recommended dilutions for the primary and secondary antibodies were used in incubations. Blots were imaged using Bio-Rad imager.

### Functional Blocking Experiment

MDA-MB-231 E-cad- and E-cad+ cells were plated in 96 well glass bottom dish at 5,000 cells/well. Wells were coated with collagen (50µg/mL) prior to seeding the cells. HECD- 1 (Invitrogen, 13-1700) was used to functionally block E-cad in MDA-MB-231+Ecad cells. The control and treated wells were imaged using live cell microscopy (Nikon Ti2) to track their migration patterns for the 4 h incubation. After the 4 h migration study, cells were rinsed 1x with DBPBS and fixed with 4% PFA for 20 min at RT. After fixation, cells were stained for various primary antibodies (Table 3). Wells were prepared for imaging as described above and images were captured using a Nikon A1 confocal and 40x oil objective. Immunofluorescence analysis was conducted by a CellProfilerTM pipeline. Briefly, nuclei and their corresponding cell bodies were masked from the DAPI and Phalloidin Channels respectively. These masks were applied to the Biomarker channel to elicit spatial expression at single-cell resolution.

### Mouse models

All mouse experiments were carried out according to protocols approved by the Johns Hopkins University Animal Care and Use Committee in accordance with the NIH Guide for the Care and Use of Laboratory Animals. All mice were housed at a temperature of 25°C under a 12 h dark/light cycle.

For the bilateral studies: Primary tumors were established with cell lines obtained from ATCC. Cells were cultured in 5% CO_2_ and 37°C. Cells were rinsed with PBS 3 times, pelleted, and resuspended in 1:1 DPBS/Matrigel (Corning). 5-week-old NSG (NOD SCID Gamma) female mice were obtained. Hair was removed from mice to ensure accurate orthotopic injection into the second mammary fat pad. Mice were briefly anesthetized using isoflurane to immobilize for the injection of 1 million tumor cells in 100 µL. For the bilateral studies, E-cadherin positive cell lines were implanted on the right side of the mouse and E-cadherin negative cell lines were implanted on the left side of the mouse.

For inhibitor studies: MDA-MB-231+E-cadherin primary tumors were established in the second mammary fat pad on the left side of 5-week-old NSG female mice. The MEK1/2 inhibitor PD0325901 (Selleckchem) was orally administered via peanut butter pellets daily for 5 days starting on day 8 post tumor cell injection at a dosage of 20 mg/kg and ending on day 12. Control group was given vehicle control (DMSO) in peanut butter pellet.

For both studies: Tumors were measured in two dimensions (x and y) with calipers every three days and weights of animals were recorded. Tumors were categorized as spheres or ellipses by calculating the difference between x and y. If x-y < 1mm, the tumor volume was calculated using the volume formula for a sphere. If x-y > 1mm, the tumor volume was calculated using the volume formula for an ellipse. At conclusion of the studies, mice were sacrificed using isoflurane and cervical dislocation. For bilateral tumor study, tumors were excised, weighed, and preserved for future studies. For inhibitor study, lungs and livers were also excised and preserved for future studies. All IHC staining was performed by an internal core facility.

For tail vein “forced metastasis” model: 5-week-old female NSG mice were placed under heat lamp for 5-7 min to dilate tail veins. 100 µL of either E-cad+ or E-cad- cell solution (5,000 cells/µL) was injected into the lateral tail vein using a 26-gauge needle. Mice were monitored every three days and sacrificed four weeks post injection using isoflurane and cervical dislocation. Lungs were harvested and then inflated with 1.5% agarose.

### Processing and analysis of harvested mouse tissues

Portions of primary tumors, lungs and liver were snap frozen in liquid nitrogen in addition to sections that were fixed in formalin were sent for sectioning and staining. H&E slides were obtained for visualization of the tumor microenvironment and regular slides were obtained for immunohistochemistry staining. Protein was extracted from the primary tumor samples using 2X SDS sample buffer. Tumor tissue was broken down using tissue grinder, samples were then heated at 100°C and sonicated. DNA was extracted from tumor tissue using DNA all prep kit (Qiagen).

### Proteomics Sample Preparation

Sample preparation for proteomic and phosphoproteomic characterization were carried out as previously described^3, 4^. In brief, tumors were excised and immediately snap frozen in liquid nitrogen to ensure maximum preservation of phosphorylated proteins.

Frozen tumor tissue was cryopulverized and resuspended in lysis buffer (8M urea, 75 mM NaCl, 50 mM Tris-HCl, pH 8.0, 1 mM EDTA, 2 µg/mL aprotinin, 10 µg/mL leupeptin, 1 mM PMSF, 10 mM NaF, Phosphatase Inhibitor Cocktail 2 and Phosphatase Inhibitor Cocktail 3 (1:100 dilution), and 20 µM PUGNAc). To remove any particulates, lysates were centrifuged at 20,000 x g for 10 min at 4°C. Protein lysates were subjected to reduction with 5mM, 1,4-Dithiothreitol for 30 min at RT. Then were subjected to alkylation with 10mM iodoacetamide for 45 min at RT, away from light. 50 mM Tris-HCl (pH 8.0) was used to reduce urea concentration to < 2M. Samples were subjected to digestion of Lys-C (Wako Chemicals) (1:50) for 2 h at RT and then to trypsin (Promega) (1:50) overnight at RT. Generated peptides were then acidified to final concentration of 1% formic acid, subjected to clean-up using C-18 SepPak columns (Waters), and then dried utilizing a SpeedVac (Thermo Scientific). Desalted peptides were then labeled with 10-plex Tandem-mass-tag (TMT) reagents following manufacturer’s instructions (Thermo Scientific). The samples were then reconstituted in volume of 20 mM ammonium formate (pH 10) and 2% acetonitrile (ACN) and subjected to basic reverse-phase chromatography with Solvent A (2% ACN, 5 mM ammonium formate, pH 10) and non-linear gradient of Solvent B (90% ACN, 5mN ammonium formatem pH 10) at 1 mL/min in the following sequence: 0% Solvent B for 9 min, 6% solvent B for 4 min, 6% increase to 28.5% solvent B over 50 min, 28% increase to 34% solvent B over 5.5 min, 34% increase to 60% solvent B over 13 min, ending with hold at 60% solvent B for 8.5 min – conducted on Agilent 4.6 mm x 250 mm RP Zorbax 300 A Extend-C18 column with 3.5 µm beads (Agilent). Collected fractions were concatenated and 5% of fractions were alqiuopted for global proteomic analysis. Samples for global proteomic analysis were then dried down and resuspended in 3% ACN and 0.1% formic acid before ESI-LC-MS/MS analysis. The remaining 95% of sample was utilized for phosphopeptide enrichment and further concatenated using immobilized metal affinity chromatrography (IMAC). Ni-NTA agarose beads were used to prepare Fe^3+^-NTA agarose beads, and then 300 µg of peptides reconstituted in 80% ACN/0.1% trifluoroacetic acid were incubated with 10 µL of Fe^3+^-IMAC beads for 30 min. Samples were then centrifuged and supernatant was removed. Beads were washed 2x and then loaded onto equilibrated C-18 Stage Tips with 80% ACN, 0.1% trifluoroacetic acid. Tips were rinsed 2x with 1% formic acid. Sample was then eluted (3x) off the Fe^3+^-IMAC beads and onto C-18 Stage tips with 70 µL of 500 mM dibasic potassium phosphate (pH 7.0). C-18 Stage Tips were washed 2x with 1% formic acid, followed by elution (2x) of phosphopeptides from C-18 stage tips with 50% ACN and 0.1% formic acid.

Following elution, samples were dried and resuspended in 3% ACN, 0.1% formic acid prior to ESI-LC-MS/MS analysis. 1 µg of peptide was separated using Easy nLC 1200 UHPLC system on an in-house packed 20 cm x 75 µm diameter C18 column (1.9 µm Reprosil-Pur C18-AQ beads (Dr. Maisch GmbH); Picofrit 10 µm opening (New Objective)). Column parameters: 50°C, 0.300 µL/min with 0.1% formic acid, 2% acetonitrile in water and 0.1% formic acid, 90% acetonitrile. Using a 6 to 30% gradient over 84 mins peptides were separated and analyzed using Thermo Fusion Lumos mass spectrometer (Thermo Scientific). Following parameters were used- MS1: 60,000 for resolution, 350 to 1800 m/z for mass range, 30% for RF Lens, 4.0e^5^ for AGC Target, 50 ms for Max IT, 2 to 6 for charge state include, 45 s for dynamic exclusion, top 20 ions selected for MS2. For MS2: 50,000 for resolution, 37 for high-energy collision dissociation activation energy (HCD), 0.7 for isolation width (m/z), 2.0e^5^ for AGC Target, and 105 ms for Max-IT.

### Proteomics Data Analysis

All LC-MS/MS files were analyzed by MS-PyCloud, a cloud-based proteomic pipeline developed at Johns Hopkins University. MS-PyCloud was used to perform database search for spectrum assignments^5^, using MS-GF+ against a combined human and mouse UniprotKB Swiss-Prot database^6, 7^. We focused on human-derived proteins for downstream analyses. False discovery rate at PSM, peptide, and protein levels were obtained using a decoy database^8^. Peptides were searched with two trypic ends, allowing for up to two missed cleavages. The following search parameters were used: 20 ppm precursor tolerance and 0.06 Da fragment ion tolerance, static modification of carbamidomethylation at cysteine (+57.02146), TMT-labeled modification of N-terminus and lysine (+229.1629) and variable modifications of oxidation of methionine (+15.99491), phosphorylation at serine, threonine, tyrosine (+79.96633). The following filters were used for global data analysis: one PSM per peptide, two peptides per protein, 1% FDR at protein level. The following filters were used for phosphoproteome data: one PSM per peptide, one peptide per protein, 1% FDR threshold at peptide level. Phosphopeptide-level data was used to examine overall relationship between substrates and kinases. Kinase-substrate associations wre extracted from PhosphoSitePlus (Hornbeck 2015) to eliminate phosphopeptides containing phosphosites that were not reported as well as those without associated kinases identified in our global dataset. We then calculated the fold changed (log_2_ scale) and then ranked each tumor (greater than 1.5 fold increase)^3, 4^.

### Generation of heatmaps and volcano plots for proteomics data

Normalized log_2_ transformed protein expression values were used to generate heatmaps. Unsupervised hierarchical clustering was used to group samples (top dendrogram) and proteins (side dendrogram) using Euclidean distances. Heatmaps were generated in R, using the pheatmap package^9^. Normalized log_2_ transformed protein expression values were used to generate volcano plots. Log_2_ fold changes were calculated between groups of samples. A two-tailed t-test was used to generate P-values. P-values were adjusted via a Benjamini-Hochberg procedure to correct for multiple comparisons. Statistically significant differential expression was determined to be greater than 1.5-fold change and an adjusted P-value < 0.05. Volcano plots were generated in R using the EnhancedVolcano package^10^.

### Proximity Ligation Assay

MDA-MB-231 E-cad- and E-cad+ cells were plated in 96 well glass bottom dish at 5,000 cells/well. Wells were coated with collagen (50µg/mL) prior to seeding the cells. After overnight incubation, cells were fixed using 4% PFA for 20 min at RT. MDA-MB-231 E- cad- cells were stained with EGFR (Rb) and E-cad (Ms) antibodies (full info in Table 3). MDA-MB-231 E-cad+ cells were stained with: EGFR (Rb) and E-cad (Ms), just EGFR (Rb), just E-cad (Ms), and with no antibodies. Duolink proximity ligation assay (Millipore- Sigma) was used to determine if EGFR and E-cad were interacting in MDA-MB-231 E-cad+ cells. After completion of the assay, wells were stained with phalloidin (f-actin) for 15 minutes at RT. Wells were then washed and DuoLink *inSitu* Mounting medium with DAPI (Millipore Sigma) was added to the wells. Confocal images were captured using a 40x oil objective and Nikon A1 Confocal microscope. The puncta in the images is first detected and counted used the previously developed particle tracking algorithm^11^. The cell and nuclei masks of individual cells were then obtained from the DAPI and phalloidin images based on the previously established segmentation algorithm^12^ and the cell masks was then used to determine the number of puncta per cell.

### Generation of Tunable E-cadherin-zsGreen Expressing Cell Lines

A DNA plasmid encoding the chemically tunable form of E-cadherin-zsGreen was stably transfected into MDA-MB-231 triple negative breast cancer cells as previously described in^13^.

### *Ex ovo* chick embryo chorioallantoic membrane (CAM) platform

Fertilized white leghorn eggs were used to generate *ex ovo* chick embryos for extravasation efficiency assays^14, 15^ and experimental metastasis assays^16^. In summary, freshly fertilized eggs (Day 0) were incubated in a Sportsman Rocker Incubator at 37°C and a relative humidity >45% for 4 days. Egg contents were removed from the eggshell as described in^14, 17^ and placed into individual weigh boats on Day 4. Each *ex ovo* embryo was returned to the incubator until Day 9 of embryonic development.

### Experimental Metastasis Assay in the CAM of Chick Embryos

Each cell line was harvested and diluted into D-PBS (Gibco Inc.) at a concentration of 2×10^5^ cells/mL. Using Day 9 chick embryos, 2×10^4^ cells (100 µL delivery volume) were intravenously (I.V.) injected into a small microvessel (20 µm diameter) to minimize bleeding post-IV injection. Equal distribution of cancer cells arrested throughout the CAM was confirmed using a fluorescence stereoscope (Nikon Inc.). Injected CAMs were returned to the incubator. At Day 18, each CAM was enumerated for the number of metastases (single metastasis = >5 cells/colony) using a fluorescence stereoscope. For the MDA-MB-231 cell line expressing the E-cadherin-zsGreen-DD protein (chemically tunable E-cadherin with Shield-1 ligand), 50 µL of Shield-1 (5 µM, 5% EtOH in D-PBS, Takeda Inc.) was I.V. injected into each embryo on Days 12, 14, and 16 as described by Leong *et al* ^13^.

### Extravasation Efficiency Assay in the CAM of Chick Embryos

Each cell line was harvested and diluted into D-PBS (Gibco Inc.) at a concentration of 1×10^6^ cells/mL. Using Day 13 chick embryos, 1×10^5^ cells (100 µL delivery volume) were intravenously (I.V.) injected into a small microvessel (20 µm diameter) to minimize bleeding post-IV injection. At t=0 after I.V. injection of cancer cells, individual cancer cells within a 1.5×1.5 cm field of view (as described in Leong *et al* ^14^) were enumerated with a fluorescence stereomicroscope and then returned to an incubator. At t=24 hours post-I.V. injection, l*ens culinaris* agglutinin-Rhodamine (Vector Laboratories Inc.) was I.V. injected (20 µg/embryo) to visualize the CAM microvasculature and vessel lumen.

After agglutinin injection, the same field of view on each embryo was enumerated for extravasated cancer cells using an upright confocal microscope (Fast A1R, Nikon Inc.). Only cells that had extravasated past the CAM, i.e., not overlapping with rhodamine signal, were enumerated.

### Statistical Analysis

Figure legends describe statistical tests used for individual experiments and also include p values.

## Proteomics Data Availability

The mass spectrometry proteomics data have been deposited to the ProteomeXchange Consortium via the PRIDE [1] partner repository with the dataset identifier PXD021545 and 10.6019/PXD021545. Please use the following username and password to access the data:

**Password:** MYmOhJvc

## Acknowledgements

We thank all members of the Wirtz Lab for discussions and feedback on this project. We specifically thank Pratik Kumat for his assistance with running the image analysis. We also thank Alan Meeker and Sujayita Roy from the Oncology Tissue Services IHC Core at Johns Hopkins Medical Campus for their assistance. This work was supported through grants from the National Cancer Institute (U54CA143868) and the National Institite on Aging (U01AG060903) to D.W. and P.H.W.

## Contributions

G.C.R. and D.W. developed the hypothesis and designed experiments. G.C.R. and M.H.L performed most experiments and data analysis. A.J.C., J.C., R.C., M.N.K., B.S. assisted with the experiments. V.W.R. assisted with proximity ligation assay. D.C., T.L., H.Z. performed proteomics experiments, assisted in the analysis, and edited the manuscript. D.J.D. and H.S.L. performed CAM experiments, assisted in the analysis, and edited the manuscript. P.W. assisted with image analysis. Q.Z. assisted with the analysis of the clinical data. G.C.R. and D.W. wrote the manuscript with input from A.J.C and V. W.R.

