## Supplementary figures and images for "E-cadherin interacts with EGFR resulting in hyper-activation of ERK in multiple models of breast cancer"

### Supplemental Figure 1

#### Extended Data Figure 1

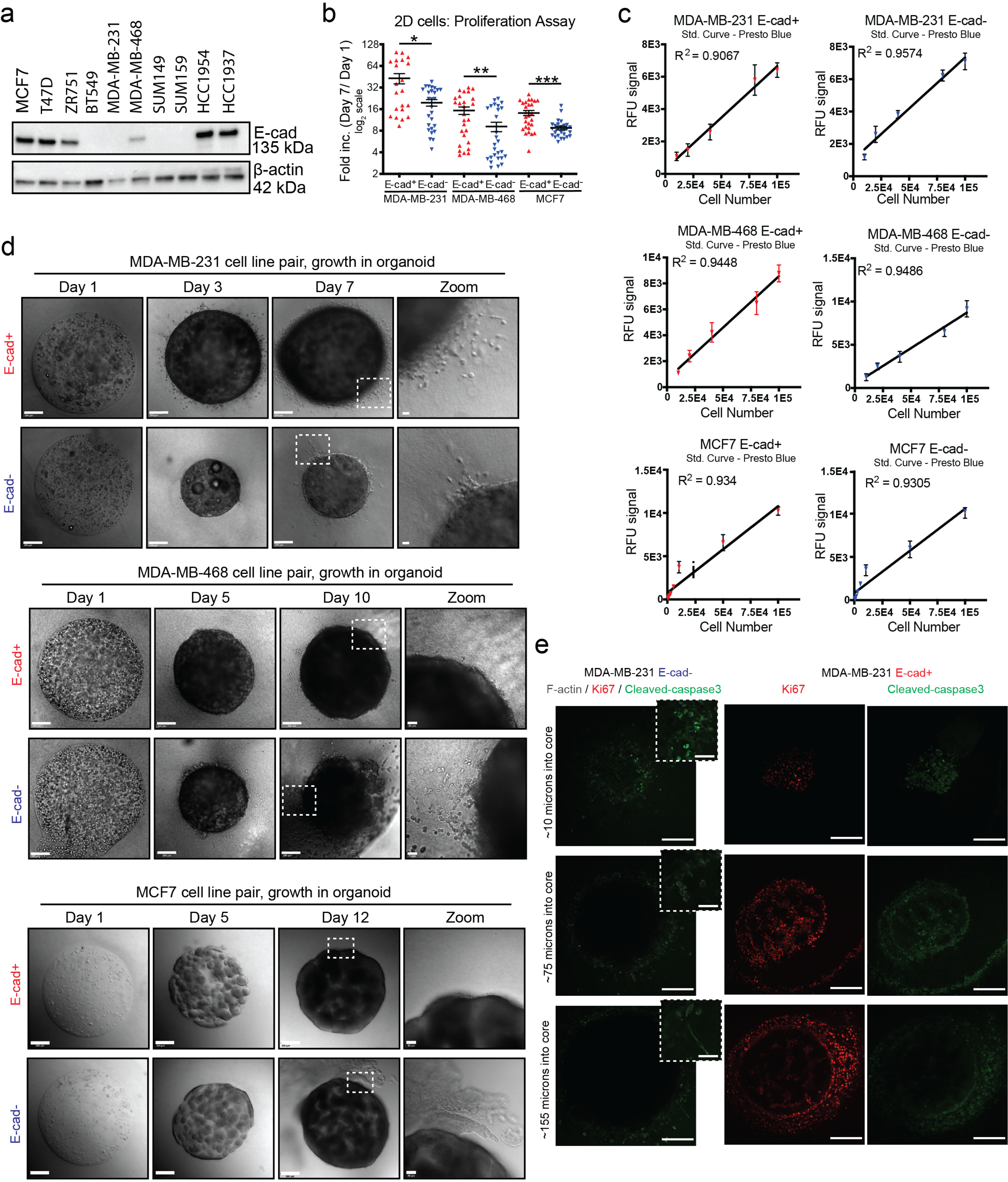

### Supplemental Figure 2

#### Extended Data Figure 2

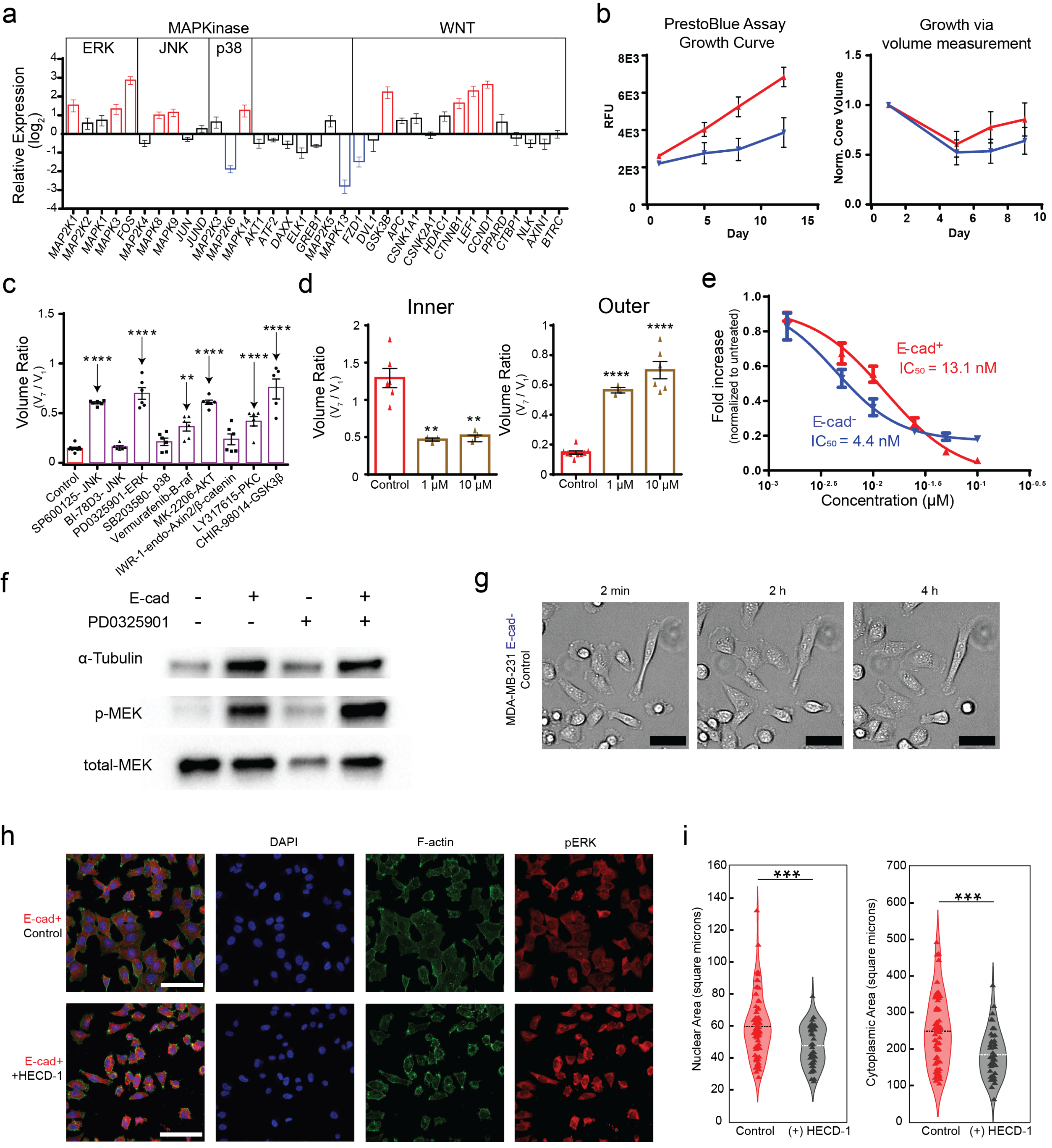

### Supplemental Figure 3

#### Extended Data Figure 3

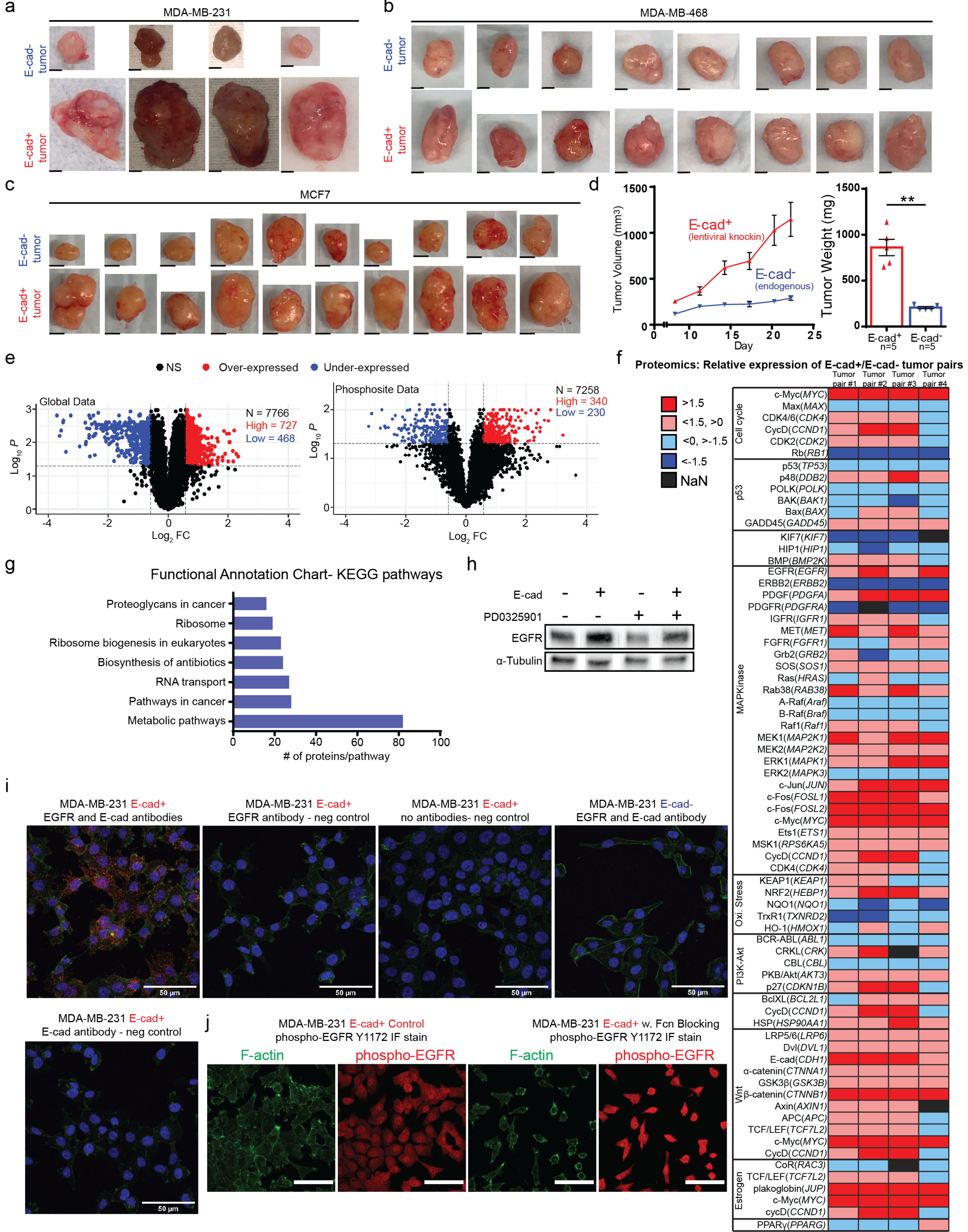

### Supplemental Figure 4

#### Extended Data Figure 4

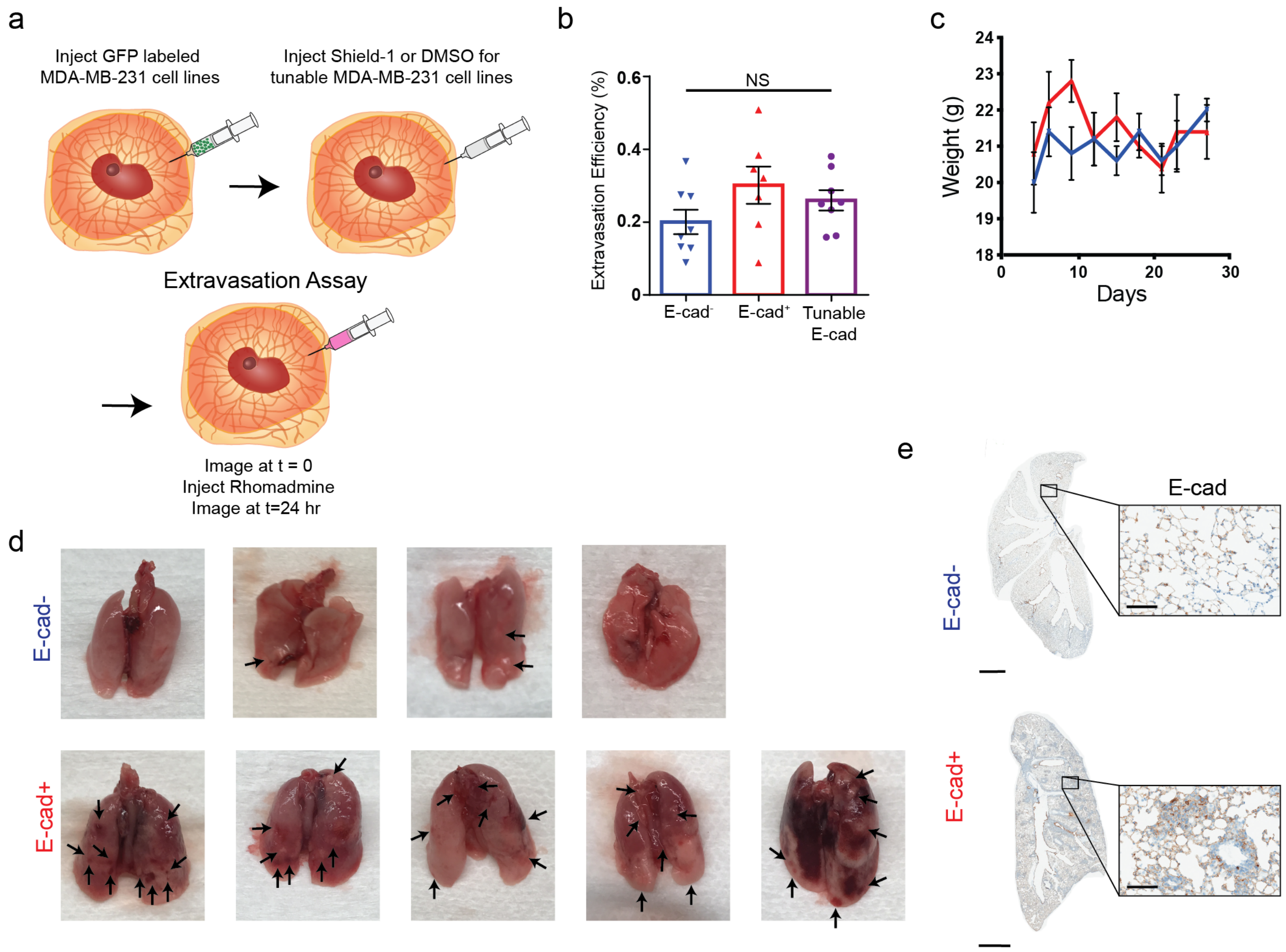

### Supplemental Figure 5

#### Extended Data Figure 5

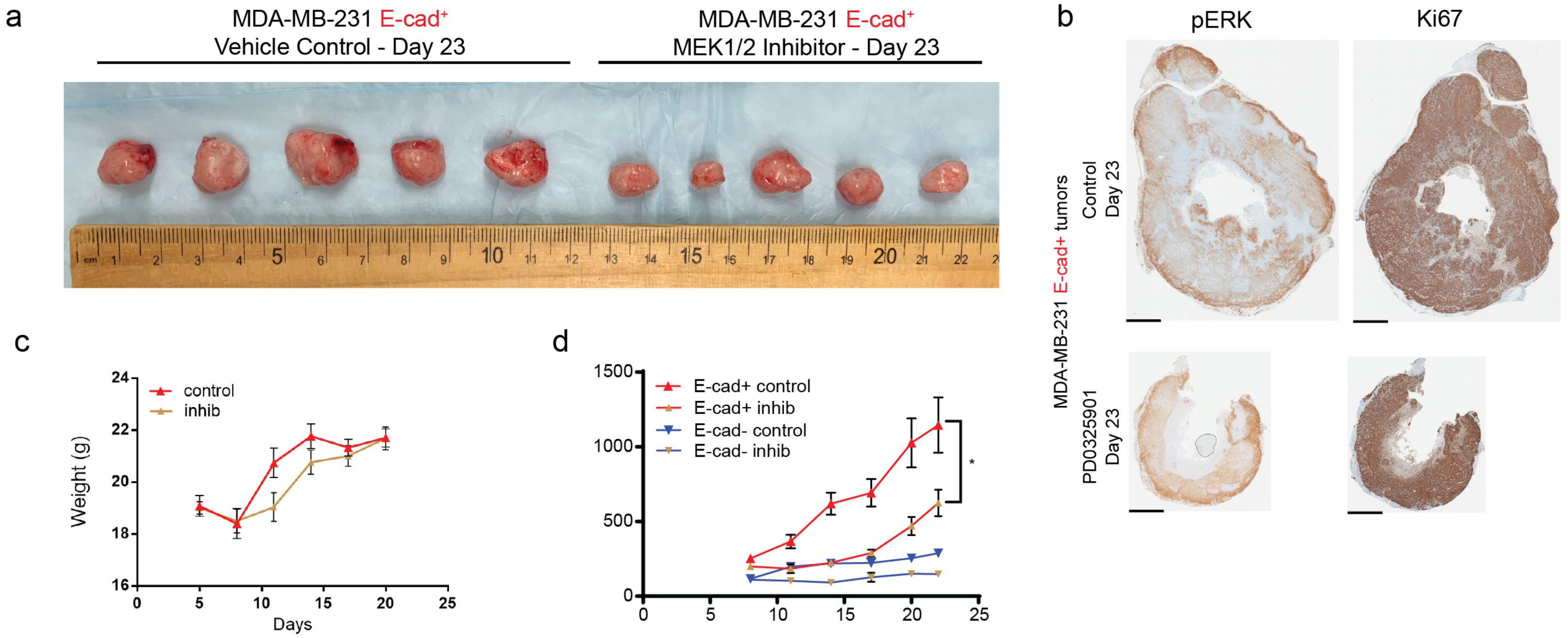
